# Transcriptome-wide *in vivo* mapping of cleavage sites for the compact cyanobacterial ribonuclease E reveals insights into its function and substrate recognition

**DOI:** 10.1101/2021.07.27.453982

**Authors:** Ute A. Hoffmann, Florian Heyl, Said N. Rogh, Thomas Wallner, Rolf Backofen, Wolfgang R. Hess, Claudia Steglich, Annegret Wilde

## Abstract

Ribonucleases are crucial enzymes in RNA metabolism and post-transcriptional regulatory processes in bacteria. Cyanobacteria encode the two essential ribonucleases RNase E and RNase J. Cyanobacterial RNase E is shorter than homologues in other groups of bacteria and lacks both the chloroplast-specific N-terminal extension as well as the C-terminal domain typical for RNase E of enterobacteria. In order to investigate the function of RNase E in the model cyanobacterium *Synechocystis* sp. PCC 6803, we engineered a temperature-sensitive RNase E mutant by introducing two site-specific mutations, I65F and spontaneously occurring V94A. This enabled us to perform RNA-seq after the transient inactivation of RNase E by a temperature shift (TIER-seq) and to map 1,472 RNase-E-dependent cleavage sites. We inferred a dominating cleavage signature consisting of an adenine at the -3 and a uridine at the +2 position within a single-stranded segment of the RNA. The data identified putative RNase-E-dependent instances of operon discoordination, mRNAs likely regulated jointly by RNase E and an sRNA, potential 3’ end-derived sRNAs and a dual-acting mechanism for the glutamine riboswitch. Our findings substantiate the pivotal role of RNase E in post-transcriptional regulation and suggest the redundant or concerted action of RNase E and RNase J in cyanobacteria.

## INTRODUCTION

Ribonucleases (RNases) play central roles in bacterial RNA metabolism (1). They are crucial for RNA degradation, but also for the maturation of mRNAs, rRNAs, tRNAs and sRNAs. Furthermore, they are involved in post-transcriptional regulation and the acclimation to changing environmental conditions, e.g. by facilitating the action of regulatory sRNAs. In bacteria, RNA degradation is assumed to be an all-or-nothing event. It is initiated by a rate-limiting step, which comprises of either 5‘-triphosphate removal or an endonucleolytic cleavage. Both result in a 5‘-monophosphorylated (5‘-P) RNA fragment. The first step is followed by rapid degradation by RNases with a high affinity to 5‘-P RNA species, such as RNase Y, RNase J or RNase E and 3‘-to-5‘ exonucleases, such as PNPase or RNase II (2). The majority of sequenced bacterial genomes encode a homologue of at least one of the three ribonucleases RNase E, RNase Y and RNase J (3). All three of these enzymes have a higher affinity to 5‘-P RNA species than to 5‘-triphosphorylated (5‘-PPP) RNAs and are able to perform the first, rate-limiting endonucleolytic cleavage initiating RNA degradation. This makes them key players in RNA metabolism. All three RNases share a low target specificity and can partially substitute for each other, which further highlights their functional similarity (4). Besides these common principles, different bacteria usually only encode a subset of the three mentioned RNases (2). The specific combination of these three RNases and peculiarities of the respective homologues, e.g. interaction with specific adaptor proteins or intracellular localisation, shape the RNA metabolism of an organism (3).

RNase E was intensively investigated in the gammaproteobacteria *Escherichia* (*E*.) *coli* and *Salmonella enterica* (*Salmonella*). Here, the enzyme plays a central role in rRNA (5, 6), tRNA (7) and sRNA (8) maturation, the action of sRNAs and bulk RNA degradation (9, 10). RNA cleavage takes place in single-stranded, adenine and uracil-rich regions (9). A uridine located two nucleotides downstream of RNase E cleavage sites was identified as an important recognition determinant in *Salmonella (8)*. The enzyme cleaves preferentially 5‘-P RNA species, which is referred to as 5‘-sensing (11). Furthermore, RNase E can recognize target RNA by secondary structures in proximity to the respective cleavage site (12, 13) or by the presence of several single-stranded regions within a single RNA target molecule (14). In addition to catalysing RNA cleavage, enterobacterial RNase E interacts with multiple proteins and serves as scaffold for the degradosome complex via its C-terminal ∼550 amino acids domain (15). In *E. coli*, RNase E interacts with specific proteins mediating the target recognition and specificity such as Hfq (16) or the adaptor protein RapZ (17). Homologues of these proteins either do not exist in *Synechocystis* (RapZ) or lost their RNA-binding capability in case of Hfq (18).

As the only bacteria that perform oxygenic photosynthesis, cyanobacteria are of immense ecological relevance. They are considered promising for the sustainable production of chemical feedstock and biofuels and serve as easy-to-manipulate models in synthetic biology and photosynthesis research (19). Cyanobacteria are morphologically distinct from other bacteria by the presence of thylakoids, extensive intracellular membrane systems, and carboxysomes, microcompartments specialised for the fixation of CO_2_. It is unknown to what extent these ultrastructural differences correlate with differences in processes such as RNA localisation and RNA metabolism. All cyanobacteria which were sequenced so far encode both RNase E as well as RNase J, but no RNase Y homologue (2). In this study, we chose *Synechocystis* sp. PCC 6803 (in the following *Synechocystis*), a unicellular model cyanobacterium. Both RNase E as well as RNase J are essential in *Synechocystis* (20–22). Besides cyanobacteria, only alphaproteobacteria, several actinobacteria and fibrobacteres contain both RNase E and RNase J, but lack RNase Y (2).

The *Synechocystis* RNase E N-terminal region is homologous to the *E. coli* RNase E N-terminal, catalytically active half. Cyanobacterial RNase E homologues contain only a short C-terminal non-catalytic domain (15). With usually less than 700 amino acids, they are shorter than the majority of other characterised RNase E homologues, which frequently consist of more than 900 amino acids (23). Moreover, cyanobacterial RNase E homologues lack the long N-terminal extension which is typical for RNase E homologues present in plants plastids (23, 24) and certain other bacteria such as *Streptomyces* (25). The C-terminal region of cyanobacterial RNase E contains several conserved microdomains, of which a cyanobacterial-specific nonapeptide is binding the 3‘-to-5‘ exonuclease PNPase (26).

Previous studies showed that RNase E in *Synechocystis* triggers the degradation of the mRNAs of the two almost identical genes *psbA2* and *psbA3* in the dark (21), while in the light the mRNAs become protected through a unique mechanism involving the asRNAs PsbA2R and PsbA3R (27). By recruiting the sRNA PsrR1, RNase E was shown to “decapitate” the *psaL* mRNA after a shift from low to high light by cleaving off a fragment consisting of the 5’ UTR and the first seven nt of the coding sequence (CDS) (28). All these mRNAs encode central proteins of the photosynthetic apparatus. Therefore, *Synechocystis* RNase E appears to be involved in the acclimation of photosynthesis to changing light conditions, consistent with the results of a recent transcriptome analysis of an RNase E knock-down mutant (20). A surprising discovery was the identification of RNase E as the major maturation enzyme of crRNAs from the *Synechocystis* CRISPR3 array (29). Studies on single transcripts, such as *Synechocystis psbA2*, CRISPR3 or *E. coli* 9S RNA and RNAI pointed towards a similar cleavage specificity of *Synechocystis* RNase E as for the enterobacterial enzyme (15, 20, 21, 29), consistent with the finding that *Synechocystis* RNase E can rescue an *E. coli* RNase E mutant (21).

Despite the multiple evidence for an important role of RNase E in *Synechocystis*, its targetome has not been determined thus far. Here, we engineered a temperature-sensitive RNase E mutant strain and applied the ‘transiently inactivation of an essential ribonuclease followed by RNA-Seq’ (TIER-seq) approach (8) for the transcriptome-wide identification of RNase-E-dependent cleavage sites. We identified 1,472 RNase-E-dependent RNA processing events, a putative sequence motif for cleavage and substantiate the function of RNase E in crRNA maturation and the regulation of essential cellular functions such as photosynthesis.

## MATERIAL AND METHODS

### Bacterial strains and culture conditions

A motile wild-type strain of *Synechocystis* was used, which is capable of photoautotrophic, mixotrophic and chemoheterotrophic growth on glucose. It was originally obtained from S. Shestakov (Moscow State University, Russia) in 1993 and re-sequenced in 2012 (30). For culturing, BG-11 medium (31) substituted with 0.3% (w/v) sodium thiosulfate and 10 mM N-[Tris(hydroxymethyl)methyl]-2-aminoethanesulfonic acid (TES) buffer (pH 8.0) was used. Liquid cultures were grown in Erlenmeyer flasks under constant shaking (135 rpm) at 30°C and continuous white light (30 µmol photons m^-2^ s^-1^) in an incubator shaker (Multitron Pro, Infors AG, Switzerland). Plate cultures were grown on 0.75% bacto-agar BG-11 plates. Kanamycin (40 μg/mL) and chloramphenicol (10 μg/mL) were added to plate cultures for strain maintenance, but omitted during experiments.

### Spectroscopy

Whole-cell absorption spectra were measured using a Specord 250 Plus (Analytik Jena) spectrophotometer at room temperature and normalized to absorption values at 682 nm and 750 nm.

### Construction of mutant strains

Plasmid pUC19-3xFLAG-*rne* was created by assembly cloning (AQUA cloning) (32). Primer pairs P01/P02 and P03/P04 (all oligonucleotides are listed in Supplementary Table S1) were used for amplification of homologous flanks by PCR. pUC19 backbone and a kanamycin resistance cassette were amplified using primer pairs P05/P06 and P07/P08 using plasmids pUC19 and pVZ321 (33) (NCBI:AF100176.1) as templates, respectively. After assembly cloning, the resulting plasmid was introduced into wild-type *Synechocystis* (WT) by transformation. Chromosomal DNA of the resulting strain was used to amplify the *rne*-*rnhB* locus, i.e. the operon of *rne* and *rnhB*, encoding RNase E and RNase HII, respectively. The PCR product included an N-terminal 3xFLAG tag, promoter and terminator sequences (Supplementary Figure 1). During amplification, XhoI and SalI recognition sites were introduced using primers P09/P10. The PCR fragment was inserted into pJET 1.2 vector (CloneJET PCR Cloning Kit, Thermo Scientific™, Germany). Point mutations were introduced by site-directed mutagenesis (Q5 Site-Directed Mutagenesis Kit, New England BioLabs, Inc., USA) using primer pairs P11/P12 (G63S), P13/P14 (I65F) and P15/P16 (G63S, I65F). The resulting sub-cloning plasmids and the conjugative plasmid pVZ321 were cleaved with restriction enzymes XhoI and SalI (Thermo Scientific™, Germany), generating compatible restriction sites, and subsequently ligated using T4 DNA ligase (New England BioLabs, Inc., USA). Insertion direction was tested by multiplexed PCR using primers P17, P18 and P19. For further strain construction, plasmids were selected for which primer pair P18/P19 yielded an amplicon of 590 bp, while P17/P18 did not give a product. For an empty-vector control, cut pVZ321 was ligated without insert, resulting in pVZΔKm^R^. The resulting plasmids were transferred into WT by triparental mating with *E. coli* DH5α harbouring the constructed plasmid and *E. coli* J53 (NCBI:txid1144303) with the conjugative helper plasmid RP4 (NCBI:txid2503).

The plasmid used for deleting the endogenous *rne-rnhB* locus was generated by AQUA cloning (Supplementary Figure 1). To construct this plasmid, four PCR fragments were generated: Homologous flanks up- and downstream of the locus were amplified using primer pairs P20/P21 and P24/P25. A kanamycin resistance cassette and pUC19 plasmid backbone was amplified using primers P22/P23 and P26/P27. After assembly cloning, the resulting plasmid was transferred into the strains harbouring the plasmids containing the *rne-rnhB* locus. For the empty-vector control, a kanamycin resistance cassette and an N-terminal 3xFLAG tag sequence were inserted at the endogenous *rne-rnhB* locus using the plasmid pUC19-3xFLAG-*rne. Synechocystis* clones were tested for temperature sensitivity by streaking them on plates which were incubated at either 30°C or 39°C. PCR was used to verify correct construct insertion and full segregation, using primers P10/P28. The occurrence of compensatory mutations and the loss of introduced point mutations were checked by PCR and sequencing with primers P29, P30, P31, P32 and P33. A list of all strains used in this study can be found in supplementary Table S2.

### Transient inactivation of RNase E and RNA extraction

For transient inactivation of RNase E, four independent liquid cultures of WT, *rne*(WT) and *rne*(Ts) were grown at 30°C to an OD_750nm_ of 0.7 – 0.8 in 50 ml culture volume. After harvesting a subsample of 20 ml culture, the temperature of the incubator shaker was set to 39°C. Further 20 ml aliquots were collected exactly 60 min after changing the set temperature. For sampling, aliquots were collected by rapid vacuum filtration through 0.8 µm polyethersulphone filter disks (Pall, Germany). Filters were immediately transferred to 1.6 ml PGTX solution (34), vortexed, frozen in liquid nitrogen and stored at -80°C. RNA was extracted according to Pinto et al. (34) with modifications as described by Wallner et al. (35). Purified RNA was quantified using a NanoDrop ND2000 spectrophotometer (Thermo Fisher Scientific, Germany).

### Northern blot hybridization

Total RNA was separated either by electrophoresis on agarose/formaldehyde gels (1.3% (w/v) agarose, 1.85% (v/v) formaldehyde, 1x MOPS-EDTA-NaOAc-buffer) or on polyacrylamide (PAA)-urea gels (8% PAA, 8.3 M urea, 1x Tris-borate-EDTA buffer) and blotted onto Roti-Nylon plus membranes (Carl Roth, Germany). Hybridisation of the membranes with radioactively labelled probes was carried out as described (29) with the following modifications: Incubation with washing solution I (2x SSC, 0.5% SDS) was performed at 25°C. Subsequent washes with buffer II (2x SSC, 0.5% SDS) and III (0.1x SSC, 0.1% SDS) were performed at 65°C. Signals were detected by phosphorimaging on a Typhoon FLA 9500 imaging system (GE Healthcare, USA). Oligonucleotides H01 - H10 used to generate hybridisation probes are given in supplementary Table S1. Single-stranded RNA probes were produced by *in vitro* transcription of PCR fragments using the Ambion T7 polymerase maxiscript kit (Thermo Fisher Scientific, Germany) in the presence of [α-32 P]-UTP (Hartmann Analytics, Germany).

### RNA sequencing

Biological triplicates of *rne*(WT) and *rne*(Ts) at each condition were analysed by RNA sequencing. These triplicates corresponded to three and two biologically independent clones, respectively. Residual DNA was removed from samples containing each 10 µg RNA by three subsequent incubation steps with Ambion TURBO DNase (Thermo Fisher Scientific, Germany). For each step, 2 U TurboDNase was added, followed by 20 min incubation at 37°C. RNA was recovered using RNA Clean & Concentrator kits (Zymo Research, USA). RNA integrity was controlled on a Fragment Analyzer using the High Sensitivity RNA Analysis Kit (Agilent, USA). cDNA libraries were constructed and sequenced as a service provided by vertis Biotechnologie AG (Germany) according to the tagRNA-Seq protocol (36), including unique molecular identifiers (UMIs) (37). Ribosomal RNAs were depleted using in-house depletion probes. 5‘-P RNA fragments were ligated to the 5‘ Illumina TruSeq sequencing adapter carrying the sequence tag CTGAAGCT, indicating processing sites (PSS). After incubation with 5‘-phosphate-dependent exoribonuclease XRN-1 (New England BioLabs, Inc., USA), samples were treated with RNA 5‘ polyphosphatase (Lucigen). The 5‘ Illumina TruSeq sequencing adapter carrying sequence tag TAATGCGC was ligated to newly formed 5‘-P RNA ends, indicating transcriptional start sites (TSS). RNA was fragmented and an oligonucleotide adapter was ligated to the resulting 3‘ ends. First-strand cDNA synthesis was performed using M-MLV reverse transcriptase and the 3‘ adapter as primer. First-strand cDNA was purified and the 5‘ Illumina TruSeq sequencing adapter was ligated to the 3‘ end of the antisense cDNA. This was followed by PCR amplification to about 10-20 ng/µl using a high fidelity DNA polymerase. cDNA was purified using the Agencourt AMPure XP kit (Beckman Coulter Genomics). Samples were pooled in approximately equimolar amounts and the cDNA pool in the size range of 200 – 600 bp was eluted from a preparative agarose gel. The cDNA pool was sequenced with 75 bp read length on an Illumina NextSeq 500 system.

### Bioinformatic analyses

A homology model of *Synechocystis* RNase E was created with iTasser (38, 39) and analysed using Pymol Molecular Graphics System, (v2.4.0) (Schrödinger, LLC.). SyntTax (40) was used for synteny analyses. For RNA-Seq analysis, reads were uploaded to the usegalaxy.eu server and analysed utilizing the Galaxy web platform (41) after preliminary processing (Supplemental Methods 1). Several workflows were created to process the data further and can be accessed and reproduced at the following links: https://usegalaxy.eu/u/ute-hoffmann/w/tier-seqprocessing1, https://usegalaxy.eu/u/ute-hoffmann/w/tier-seqtranscript, https://usegalaxy.eu/u/ute-hoffmann/w/tier-seqpss-tss and https://usegalaxy.eu/u/ute-hoffmann/w/tiertranscript-coverage. For downstream analyses, reads with a mapping quality above 20 were kept. Reads with a mapping quality of exactly one were included for analyses of *psbA2*/*psbA3* and rRNA loci. Transcript data was assigned to annotated regions using htseq-count (42). Annotation included several known sRNAs, asRNAs and small proteins (43–46). Subsequently, htseq-count files, PSS/TSS-5‘ end files and transcript coverage files were downloaded and analysed using Python (v3.7.4) and R (v4.0.4) scripts available at (https://github.com/ute-hoffmann/TIER-synechocystis) (Supplementary Methods 2). DESeq2 (v1.30.1) (47) was used for the analysis of htseq-count files of transcript data (|log2FC| > 0.8 and p.adj < 0.05), the classification of genomic positions as TSS or PSS and differential expression analysis of the resulting set of PSS and TSS positions (|log2FC| > 1.0 and p.adj < 0.05). Sequence logos were created using WebLogo (v3.7.8) (48) with a GC content of 47.4%. Minimal folding energies (ΔG) were calculated with RNA fold (v2.4.17) (49), temperature parameter set to 39°C, with a sliding window of 25 nt and a 1 nt step size.

## RESULTS AND DISCUSSION

### A point mutation homologous to *E. coli rne-3071* conveys temperature sensitivity in the cyanobacterium *Synechocystis*

*Synechocystis* RNase E, encoded by *rne*, is essential (20–22) and forms an operon with *rnhB* (*slr1130*) encoding RNase HII (46), which is a widely conserved arrangement in cyanobacteria (Supplementary Figure 2). Analyses in *E. coli, Salmonella* and *Rhodobacter sphaeroides* (*Rhodobacter*) demonstrated that temperature-sensitive RNase E mutant strains enable the transient inactivation of the essential RNase to perform RNA-sequencing and capture RNase-E-dependent cleavage sites on a transcriptome-wide level *in vivo* (8, 14, 50). Despite the low sequence identity of 36% and similarity of 55% between *Synechocystis* RNase E and the catalytic, N-terminal half of the *E. coli* enzyme (20), we aimed for a temperature-sensitive RNase E mutant strain in *Synechocystis*. To avoid disruption of the *rne*-*rnhB* operon and to minimize a polar effect on *rnhB* expression, we first introduced the *rne*-*rnhB* locus including the native promoter and 3‘ UTR on a conjugative self-replicating plasmid into WT cells. This was followed by deletion of the genomic locus by homologous recombination (Supplementary Figure 1).

In *E. coli*, studies employing temperature-sensitive RNase E relied on one of the two point mutations *ams-1* (G66S) or *rne-3071* (L68F) (51). Using sequence alignments and a homology model, we identified G63S and I65F as the homologous amino acid substitutions in *Synechocystis* (Figure 1A, Supplementary Figure 3). The respective mutated genes were introduced into *Synechocystis*, following the same strategy as for the unmodified gene. Segregation between chromosomes with the deleted *rne* gene and those still carrying the WT *rne* gene was judged by PCR (Supplementary Figure 1B). Full segregation (homozygosity) was only obtained for the mutation homologous to *rne-3071* (*Synechocystis* I65F), but not to *ams-1* (*Synechocystis* G63S). Interestingly, all homozygous clones accumulated one of the three following second-site mutations within the RNase E gene: V94A, V297A or G281E. I65F as well as V94A and V297A are localized at the RNA-binding channel of RNase E, close to the enzyme’s active site (Supplementary Figure 3B). Phenylalanine is bulkier than isoleucine. Therefore, the I65F substitution seems to destabilize the beta sheet in which it is located (*E. coli* L68, compare (52)). Substitution of V94 or V297 by alanine reduces this steric problem (Supplementary Figure 3E). The third mutation, G281E, is not localized within the RNA-binding channel, but in close proximity. It is part of a beta sheet which forms the RNA-binding channel, but G281E is oriented towards the protein’s outer surface. How the mutation G281E might compensate for I65F is not directly apparent, since it does not seem to lower the steric issues associated with I65F. It might be part of a potential RNA-binding surface which is possibly involved in the 5‘ bypass pathway (12) (Supplementary Figure 3D). Of all three detected second-site mutations, V94A seems to have the smallest effect on potential RNA-binding sites and the RNA channel, according to the homology model. Hence, for further characterization, the I65F and V94A mutations were combined in one strain, referred to as *rne*(Ts). The congenic control strain, which was generated in the same manner as *rne*(Ts) and in which the RNase E WT allele was introduced, will be referred to as *rne*(WT). In *rne*(WT), no second-site mutations within the *rne* gene were detected. In addition, both RNase E variants were tagged with an N-terminal 3xFLAG tag. Previous studies showed that neither the tag nor expression from a conjugative vector interfere with RNase E function (29).

**Figure 1.**
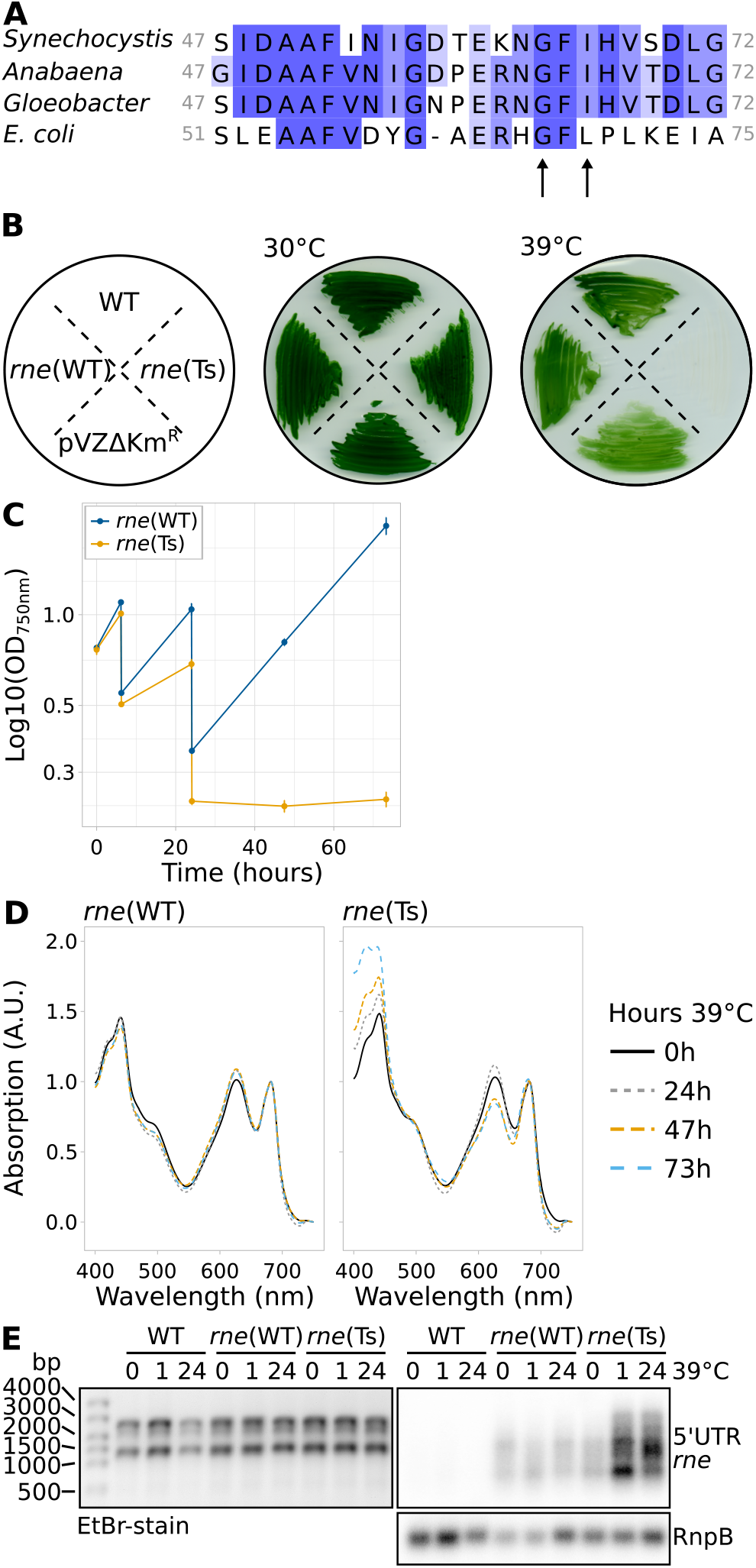
Characterisation of *Synechocystis* harbouring temperature-sensitive RNase E. (**A**) Alignment of *Synechocystis* RNase E residues 47 to 72 with the respective section in homologues from *Anabaena* sp. PCC 7120, *Gloeobacter violaceus* PCC 7421 and *E. coli*. The arrows point at two conserved residues which were mutated to obtain temperature-sensitivity. (**B**) Growth of wild-type *Synechocystis* (WT), *rne*(WT), *rne*(Ts) and an empty-vector control strain (pVZΔKm^R^) at the standard temperature of 30°C and at 39°C. The scheme on the left indicates the streaking order of the strains. (**C**) Growth of *rne*(WT) and *rne*(Ts) at 39°C in liquid culture. Time point 0 h corresponds to the switch from 30°C to 39°C. Error bars indicate the standard deviation of biologically independent duplicates. Cultures were diluted after 6 hours and 24 hours of growth. (**D**) Absorption spectra of *rne*(WT) and *rne*(Ts) throughout the course of 73 h incubation at 39°C of one representative experiment (compare panel C). Spectra were normalized to absorption at 682 nm and 750 nm. (**E**) Northern blot analysis of the accumulation of the *rne*-*rnhB* transcript. One representative analysis is shown (n=4). In addition to the control hybridization with RnpB (lower panel), the denaturing agarose gel which was used for blotting is shown on the left as a loading control.

Growth of *rne*(Ts) was not impaired in comparison to WT or *rne*(WT) at 30°C. We determined 39°C as non-permissive growth temperature at which, in contrast to *rne*(Ts), neither WT nor *rne*(WT) showed a severe growth deficiency as judged by growth on solid and liquid media (Figures 1B, 1C). Prolonged incubation of *rne*(Ts) liquid cultures at 39°C resulted in bleaching of the cells (Figure 1D).

The used mutation strategy led to an overexpression of the *rne*-*rnhB* transcript, which was likely caused by the higher copy number of the RSF1010-derived conjugative plasmid compared to the chromosome (Figure 1E) (33, 53). In *Synechocystis*, the enzymatic activity of RNase E is rate-limiting for the maturation and accumulation of crRNAs from the CRISPR3 array (29). Hence, the amount of mature CRISPR3 crRNA may be used as a proxy for RNase E activity. Indeed, overexpression of RNase E led to a dramatic overaccumulation of mature CRISPR3 spacers in *rne*(WT) and a moderately enhanced accumulation in *rne*(Ts) compared to WT (Supplementary Figure 4). With increasing time at the non-permissive temperature, the level of CRISPR3 accumulation decreased in all three compared strains.

*E. coli* RNase E autoregulates its own transcript level (54) and there is evidence that the same holds in the cyanobacteria *Synechocystis* and *Prochlorococcus* MED4 (20, 55). Here, we observed enhanced *rne-rnhB* transcript accumulation in *rne*(Ts) during prolonged incubation at 39°C (Figure 1E). This finding further supported that RNase E activity was reduced in *rne*(Ts) at the elevated temperature. We conclude that introduction of the I65F mutation homologous to *E. coli rne-3071* in combination with the V94A substitution led to a temperature-sensitive RNase E *Synechocystis* mutant strain, *rne*(Ts).

### Overview of TIER-seq experiment

We used the here generated strains *rne*(Ts) and *rne*(WT) to identify RNase-E-dependent cleavage positions on a transcriptome-wide level by performing TIER-seq (Figure 2A) (8). Inactivation of a ribonuclease should lead to increased amounts of its target RNAs. Hence, transcripts accumulating after the heat shock at a higher level in *rne*(Ts) compared to *rne*(WT) likely are direct targets of RNase E. A lowered level indicates RNA species which are normally matured by the respective RNase from an otherwise unstable precursor or may derive indirectly from the RNase E inactivation (20). The processed CRISPR3 crRNAs are an example for the former and became more abundant in *rne*(WT) compared to *rne*(Ts) and WT here (Supplementary Figure 4), consistent with previous observations (29). To further pinpoint direct targets and cleavage positions of RNase E, we analysed changes in the number of processing sites (PSS) upon RNase E inactivation. We assumed that lowered PSS counts in *rne*(Ts) compared to *rne*(WT) after transient RNase E inactivation indicated direct targets of RNase E *in vivo* or resulted from the combined action of RNase E and downstream processing by other RNases. Conversely, higher PSS counts in *rne*(Ts) after heat inactivation relative to *rne*(WT) point to RNA fragments produced by the action of other RNases and which would possibly usually be further degraded by RNase E.

**Figure 2.**
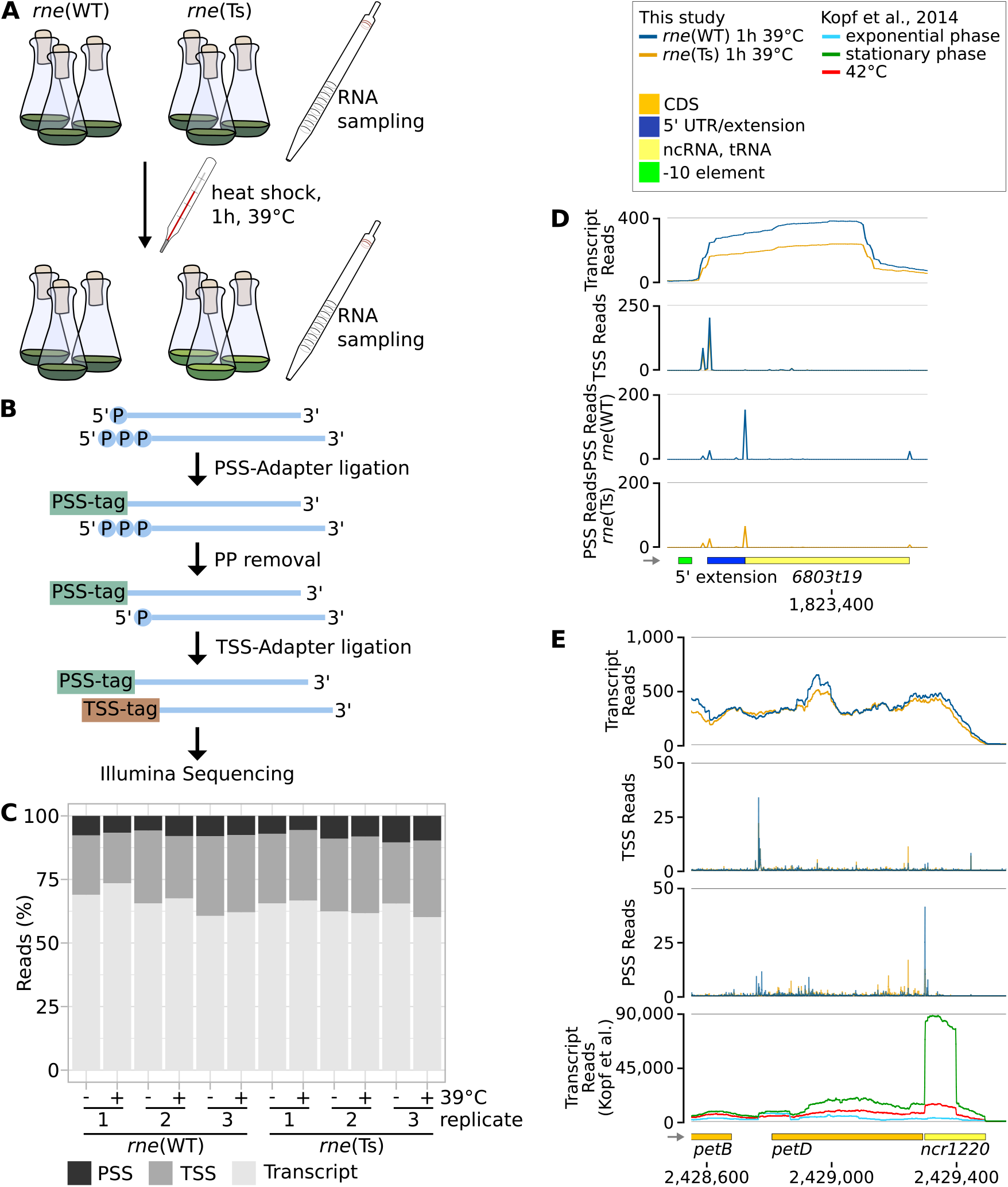
Overview of the TIER-seq experiment and exemplary results. (**A**) Experimental work flow. RNA was sampled from strains encoding either wild-typic (*rne*(WT)) or temperature-sensitive (*rne*(Ts)) RNase E before and after incubation at 39°C for one hour. (**B**) RNA was extracted and used to identify processing sites (PSS) and transcriptional start sites (TSS) after the addition of PSS- and TSS-tags, sequencing adaptors containing PSS- and TSS-specific nucleotide combinations. For the latter, 5‘ pyrophosphates were removed using RNA 5‘ polyphosphatase. (**C**) Shares in reads corresponding to PSS, TSS or unspecified transcript positions in the 12 samples. (**D**) TIER-seq data for the tRNA *6803t19*. (**E**) TIER-seq data for the CDS of *petD* and ncRNA *ncr1220*. In panels D and E, transcriptome coverage, PSS and TSS represent the average of normalised read counts of the three investigated replicates for the indicated strains. In panel E, this is compared to read counts obtained by Kopf et al. (46). CDS: coding sequences, UTR: untranslated region.

Since RNase E processing results in 5‘-P RNA ends (56), we chose a library preparation protocol that distinguishes between PSS and TSS RNA ends: tagRNA-Seq (Figure 2B) (36). Briefly, 5‘-P RNA species were ligated to a sequencing adaptor carrying a nucleotide combination specific for PSS: a PSS-tag. Unligated 5‘-P RNA fragments were removed using the 5‘-phosphate dependent exoribonuclease XRN-1. Subsequently, 5‘-PPP ends (characteristic of unprocessed TSS) were converted to 5‘-P ends and ligated to a TSS-tag followed by standard library preparation and sequencing. For sequencing, triplicates of RNA samples were taken before and after one hour of growth at 39°C of *rne*(WT) and *rne*(Ts). Samples were taken at an OD_750nm_ of 0.7 – 0.8. In total, ∼160 million raw sequencing reads were obtained for the twelve samples. After mapping, UMI-removal and quality filtering, 48.3% of those reads remained, corresponding, on average, to 5.5 million reads (∼0.4 billion bp) per sample. Of those, on average, 9.5% were PSS reads, 15.7% TSS reads and 74.5% transcript reads (Figure 2C).

### Characterization of newly identified TSS and PSS

Following the initial data analysis, 3,540 nucleotide positions were classified as TSS and 3,450 as PSS (Supplementary Methods 2, Supplementary Figure 5A, supplementary file TSS_PSS_rneAnalysis.gff). Newly identified PSS were scrutinized to assure the feasibility of our approach to identify and classify TSS and PSS. We explored the sequence composition up- and downstream of the identified TSS and PSS positions (Supplementary Figure 5B). TSS positions showed an enrichment of a canonical -10 element (5‘-TAnAAT-3‘) and a higher frequency of purines at the +1 position matching previous observations (44). PSS positions showed a slight enrichment of T 2 nt downstream, reminiscent of the motif identified by Chao et al. for *Salmonella* RNase E (8).

The majority of PSS (98.7%) overlapped with transcriptional units (TUs) defined previously (46) and coding regions of *Synechocystis*. In total, 16.0% (656) TUs and 12.0% (737) CDS contained processing sites. We noticed a correlation between the expression level and the detection of PSS (Supplementary Results 1, Supplementary Figure 5C, Supplementary Tables S3 – S6) indicating that processing sites in low abundance transcripts were possibly partially not captured.

We compared the newly obtained set of PSS to the total of 5,162 TSS determined by Kopf et al. (46). In the following, those TSS will be referred to as anno-TSS, for “annotated TSS”. Only 47 of the PSS identified using tagRNA-Seq overlapped with anno-TSS. Of these, 29 (61.7%) anno-TSS were previously classified as alternative or internal TSS within another transcript (Supplementary Figure 5D), including 11 positions which correspond to the 5‘ ends of matured tRNAs. Using tagRNA-Seq, both the 5‘-P ends of mature tRNAs as well as corresponding TSS upstream of those positions were detected (Figure 2D). Those examples show that tagRNA-Seq efficiently discriminated PSS and TSS. Some further PSS coinciding with anno-TSS were located upstream of non-coding TUs in the 3‘ UTRs of coding genes, e.g. TU2529 (*ncr1220*), which is located in the 3‘ UTR of the *petBD* operon (encoding cytochrome *b*6 and subunit 4 of the cytochrome *b*_6_*f* complex) (Figure 2E). Under certain growth conditions, e.g. in the stationary phase, TU2529 accumulates differently from the *petBD* main transcript (46). TU287, representing the 3’ UTR of *ycf19*, showed a similar behaviour.

To evaluate the effect of incubating the cells at 39°C, we analysed transcriptomic differences before and after the heat treatment (Supplementary Results 2, Supplementary Figure 6, Supplementary Tables S7 – S12). In both strains, the set of most strongly upregulated genes after 39°C treatment encoded heat shock proteins, while genes involved in the acclimation to inorganic carbon limitation were down-regulated.

### Transient inactivation of RNase E affects a high proportion of the transcriptome

The TIER-seq data set was separately analysed for the three different data types encompassing transcript patterns, TSS and PSS (Supplementary Tables S13 – S19). Principal component analyses (PCAs) indicated similar gene expression patterns and consistency between biological replicates and a stronger effect of the *rne*(Ts) mutation on processing rather than on transcription initiation or transcript patterns (Figure 3A). This is reflected by the number of differentially expressed transcripts, PSS and TSS positions before and after the heat treatment (Table 1, Supplementary Figure 7, p.adj < 0.05; for RNA features: |log_2_FC| > 0.8, for TSS/PSS: |log_2_FC| > 1). The 1,472 PSS with significant different read counts between both strains after the heat shock mapped to 380 (9.4%) of the annotated TUs (*rne*(WT): 248; *rne*(Ts): 237) and 307 (8.4%) annotated CDS (*rne*(WT): 198; *rne*(Ts): 182) (Figure 3B, Table 2). PSS corresponding to major rRNA maturation intermediates were not affected by RNase E inactivation (Supplementary Results 3). tRNAs levels were higher in *rne*(WT) than in *rne*(Ts) after RNase E inactivation (Table 3, Supplementary Figure 8A, Supplementary Tables S20). This indicates a role of *Synechocystis* RNase E in the maturation of tRNAs and is in line with known functions of enterobacterial RNase E (8, 57). Combined with the high proportion of transcripts affected by transient RNase E inactivation (15.5% of annotated transcriptional units), these findings illustrate the central role of RNase E in cyanobacterial RNA metabolism. Gene set enrichment analyses (GSEA) and functional enrichment analyses identified mainly photosynthesis-associated KEGG and GO terms among the differentially regulated transcripts and PSS with divergent read counts (Supplementary Results 4, Supplementary Figure 8B and 8C, Supplementary Tables S20 – S25).

**Figure 3.**
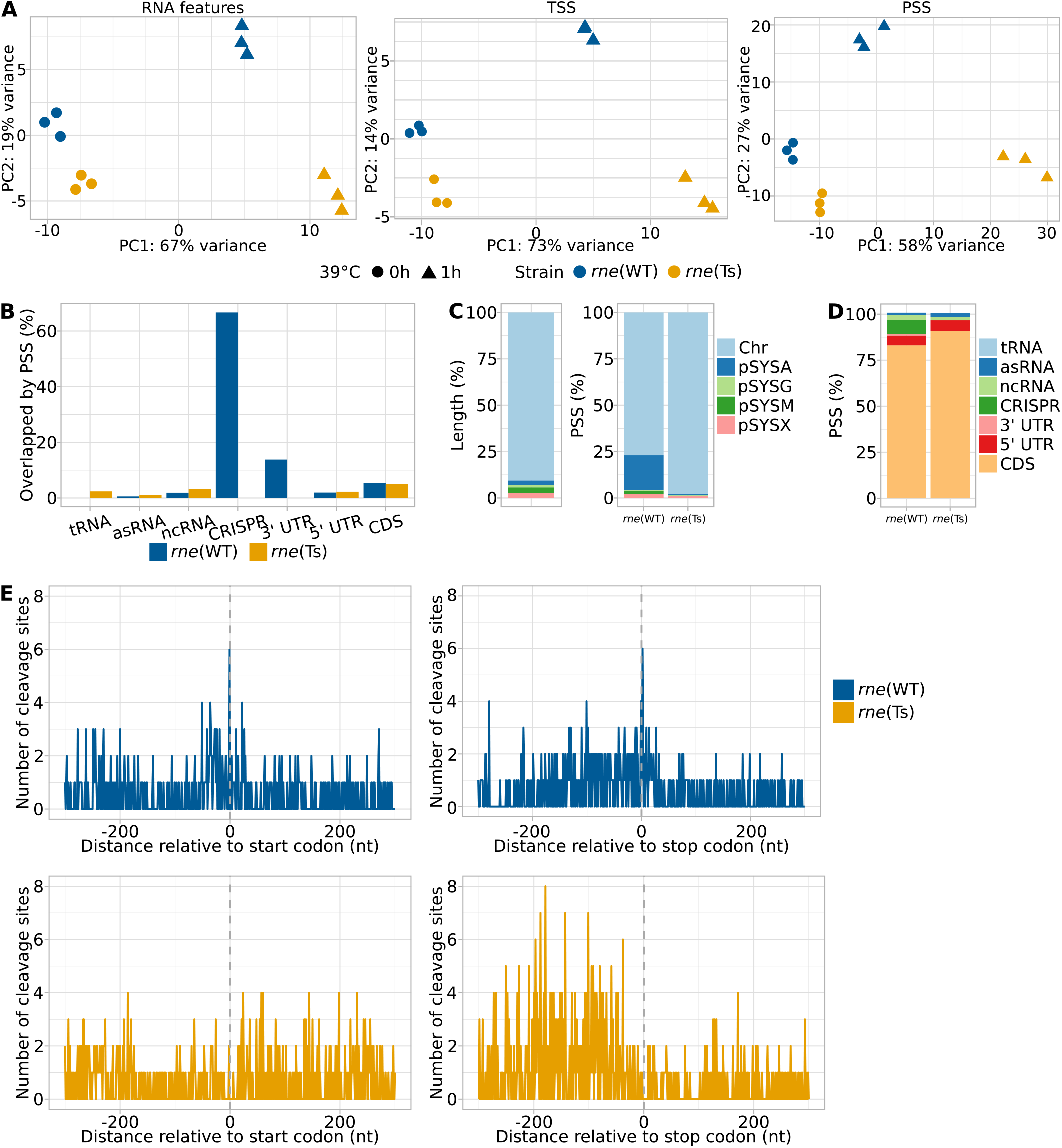
Global analysis of TIER-seq results. (**A**) Comparison between the triplicates for the three analysed strains using principal component analyses and differentiating between unspecified transcripts, TSS and PSS. (**B**) Percentages of different types of transcripts associated with PSS identified in *rne*(WT) and *rne*(Ts) after the heat treatment relative to total number of the respective transcript type in the annotation. (**C**) Relative lengths of different parts of the *Synechocystis* genome (on the left). Percentage of PSS mapped to the five major replicons comparing *rne*(WT) and *rne*(Ts) after the heat treatment (on the right). Chr: chromosome. (**D**) Percentages of PSS associated with different types of transcripts comparing *rne*(WT) and *rne*(Ts) after the heat treatment. (**E**) Distribution of PSS identified in *rne*(WT) and *rne*(Ts) after heat shock relative to start and stop codons.

**Table 1.**
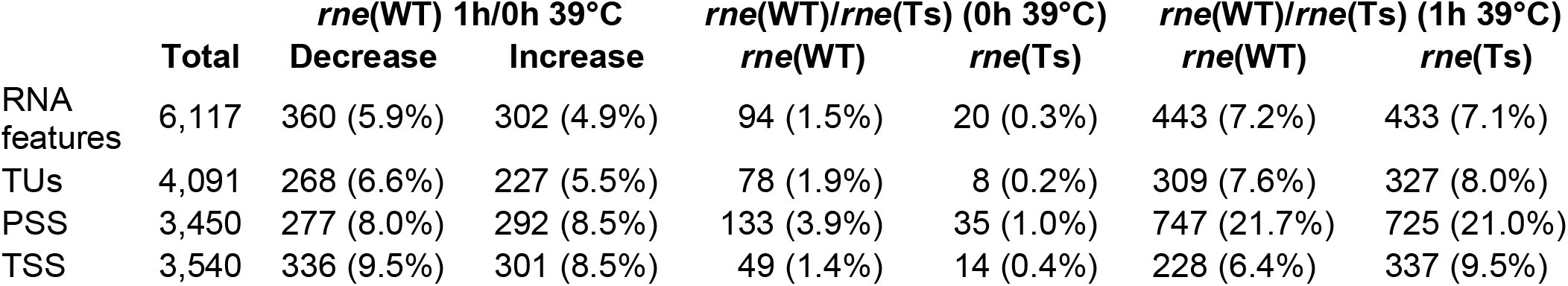
RNA features, transcriptional units (TUs), PSS and TSS differentially expressed between *rne*(WT) and *rne*(Ts) and in *rne*(WT) upon heat treatment. RNA features include CDS, known sRNAs, asRNAs and small proteins (43–45) as well as 5’ UTRs and 3’ UTR based on the total of 4,091 TUs defined by Kopf et al. (46). Numbers in parentheses give percentages relative to all features of respective type.

|  | Total | *rne*(WT) 1h/0h 39°C |  | *rne*(WT)/*rne*(Ts) (0h 39°C) |  | *rne*(WT)/*rne*(Ts) (1h 39°C) |  |
| --- | --- | --- | --- | --- | --- | --- | --- |
|  |  | Decrease | Increase | *rne*(WT) | *rne*(Ts) | *rne*(WT) | *rne*(Ts) |
| RNA features | 6,117 | 360 (5.9%) | 302 (4.9%) | 94 (1.5%) | 20 (0.3%) | 443 (7.2%) | 433 (7.1%) |
| TUs | 4,091 | 268 (6.6%) | 227 (5.5%) | 78 (1.9%) | 8 (0.2%) | 309 (7.6%) | 327 (8.0%) |
| PSS | 3,450 | 277 (8.0%) | 292 (8.5%) | 133 (3.9%) | 35 (1.0%) | 747 (21.7%) | 725 (21.0%) |
| TSS | 3,540 | 336 (9.5%) | 301 (8.5%) | 49 (1.4%) | 14 (0.4%) | 228 (6.4%) | 337 (9.5%) |

**Table 2.** Numbers of different RNA features overlapping PSS. Numbers in parentheses give percentages relative to all RNA regions of respective type.

| Type of RNA feature | Total number of RNA feature | *rne*(WT) | *rne*(Ts) |
| --- | --- | --- | --- |
| CDS | 3675 | 198 (5.4%) | 182 (5.0%) |
| 5UTR | 979 | 19 (1.9%) | 22 (2.3%) |
| 3UTR | 29 | 4 (13.8%) | 0 (0.0%) |
| tRNA | 42 | 0 (0.0%) | 1 (2.4%) |
| rRNA | 6 | 0 (0.0%) | 0 (0.0%) |
| ncRNA | 318 | 6 (1.9%) | 10 (3.1%) |
| asRNA | 1071 | 6 (0.6%) | 11 (1.0%) |
| CRISPR | 3 | 2 (66.7%) | 0 (0.0%) |

**Table 3.** Number of RNA features differentially expressed between *rne*(WT) and *rne*(Ts) after heat treatment and in *rne*(WT) upon heat treatment. Numbers in parentheses give percentages relative to all RNA regions of respective type.

|  | **Total** | ***rne*(WT) 1h/0h 39°C** |  | ***rne*(WT)/*rne*(Ts) (1h 39°C)** |  |
| --- | --- | --- | --- | --- | --- |
|  |  | **Decrease** | **Increase** | ***rne*(WT)** | ***rne*(Ts)** |
| CDS | 3,675 | 292 (7.9%) | 193 (5.3%) | 333 (9.1%) | 226 (6.1%) |
| 5' UTR | 979 | 26 (2.7%) | 25 (2.6%) | 29 (3.0%) | 41 (4.2%) |
| 3' UTR | 29 | 2 (6.9%) | 3 (10.3%) | 2 (6.9%) | 1 (3.4%) |
| tRNA | 42 | 5 (11.9%) | 0 (0.0%) | 17 (40.5%) | 0 (0.0%) |
| asRNA | 1,071 | 24 (2.2%) | 52 (4.9%) | 31 (2.9%) | 140 (13.1%) |
| ncRNA | 318 | 10 (3.1%) | 28 (8.8%) | 28 (8.8%) | 24 (7.5%) |
| CRISPR | 3 | 0 (0.0%) | 0 (0.0%) | 3 (100.0%) | 0 (0.0%) |

### RNase E inactivation affects transcripts originating from plasmid pSYSA

In relation to the sizes of the chromosome and the seven different native plasmids, PSS located on pSYSA were overrepresented among PSS accumulating in *rne*(WT) compared to *rne*(Ts) after RNase E heat inactivation (Figure 3C). This is also reflected by the higher percentage of annotated transcripts containing RNase-E-dependent PSS and the higher number of PSS per kb (Supplementary Figure 9). A large proportion of the mapped PSS correspond to processing events in the CRISPR3 array, consistent with previous observations that RNase E is the major maturation enzyme of the pSYSA-located CRISPR3 array, a type III-Bv CRISPR-Cas system (29) (Figure 4A). However, we also noticed PSS in the type I-D CRISPR1 system for which Cas6-1 was identified as the main maturation endonuclease, that cleaves 8 nt from the ends of the repeats (58). Conversely, the here identified RNase E PSS are located close to the 5’ ends of the CRISPR1 repeats suggesting possible involvement of this enzyme in the maturation or degradation of additional CRISPR RNAs. Additionally, all three CRISPR arrays were more highly expressed in *rne*(WT) than in *rne*(Ts) after the heat shock (Table 3). The large number of remaining pSYSA-located PSS might indicate further roles of RNase E associated with the multiple toxin-antitoxin systems on this plasmid (59).

**Figure 4.**
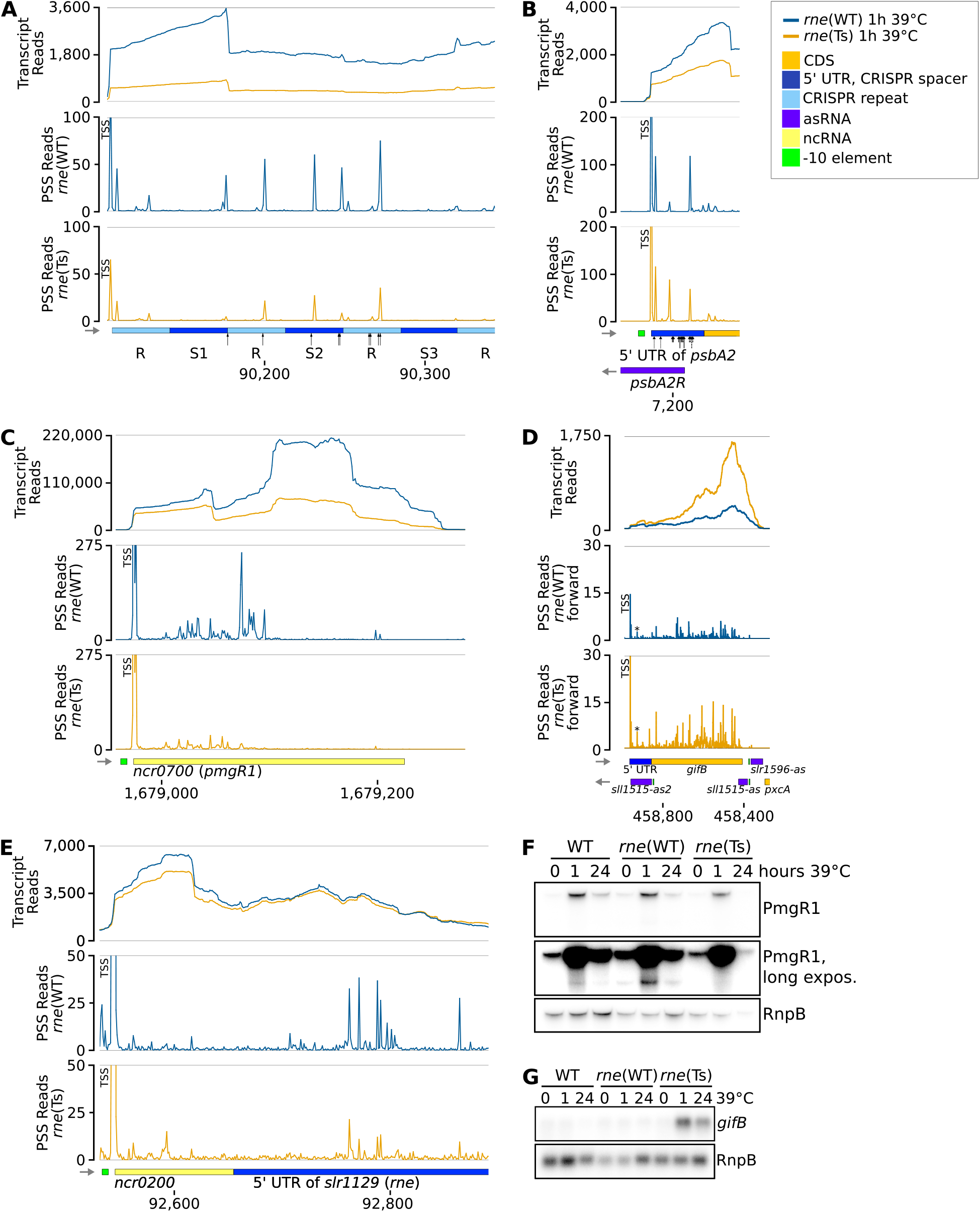
Examples of RNase-E-dependent processing events. (**A**) Spacers 1, 2 and 3 (S1 – S3) of the CRISPR3 array. The black arrows point at positions at which cleavage events were mapped previously by primer extension (29). (**B**) The 5’ UTR and begin of the *psbA2* (*slr1311*) CDS and associated asRNA PsbA2R. Black arrows point at major (long arrows) and minor (short arrows) cleavage sites detected by *in vitro* cleavage assays previously (20, 21, 29). (**C**) The sRNA PmgR1 (Ncr0700). (**D**) The CDS of *gifB* (*sll1515*), its 5’ UTR harbouring the glutamine riboswitch and two asRNAs. The prominent PSS at position +35 of the *gifB* transcript is labelled by an asterisk. (**E**) The 5‘ UTR of the *rne* gene (*slr1129*). (**F**) Representative northern blot analysis of PmgR1 (n=4). A longer exposure is shown to highlight low abundance processing products. (**G**) Representative northern blot analysis of *gifB* (n=4). The transcriptome coverage is given on top for the two indicated strains after incubation for 1 h at 39°C. Cleavage sites are displayed in the diagrams underneath by the blue and orange peaks, representing 5‘-P (PSS) detected in *rne*(WT) and *rne*(Ts), respectively. 5’-PPP (TSS) may be converted to 5’-P RNA ends *in vivo* and also during RNA-seq library preparation. Hence, TSS show up in the PSS signal. Positions which were classified as TSS using DESeq2 are labelled with “TSS” next to the respective peaks. Transcriptome coverage and cleavage sites (PSS) represent the average of normalised read counts of the three investigated replicates. CDS: coding sequences, UTR: untranslated region.

### RNase-E-dependent cleavage sites tend to be close to start and stop codons

To further analyse specific effects of RNase E, we evaluated which type of RNA features overlapped with the detected differentially accumulating PSS (Figure 3D, Table 3). The majority of PSS were mapped to CDS in both *rne*(WT) as well as *rne*(Ts). PSS mapping to 3‘ UTRs were only detected in *rne*(WT), but not in *rne*(Ts) (Figure 3B and 3D). Also, we observed a tendency for 3‘ UTRs to be enriched in *rne*(WT) compared to *rne*(Ts) after the heat shock (Supplementary Figure 8A, Supplementary Table S20).

The evidence towards an action of RNase E on 3‘ UTRs led us to a transcriptome-wide investigation of PSS localisation in start and stop codon regions (Figure 3E). Indeed, PSS accumulating in *rne*(Ts) had a slight preference to be just in the region upstream of stop codons. Intriguingly, several PSS accumulating in *rne*(WT) were located exactly 1 nt upstream of start codons or 2 nt downstream of stop codons. A similar relation was previously observed for *Salmonella* (8) and regarding start codons for *Rhodobacter* (50).

RNase-E-dependent cleavage sites 2 nt downstream of stop codons were identified for *psaJ* (*sml0008*, photosystem I subunit IX), *psbO* (*sll0427*, photosystem II manganese-stabilizing polypeptide), *rps14* (*slr0628*, 30S ribosomal protein S14) and three genes of unknown function: *ssr6030, slr5110* and *slr0581*. Indeed, the 3’ UTRs of *psaJ, psbO* and *rps14* were previously identified to accumulate different from their CDS under certain growth conditions (46), This also holds for 3’ UTRs of *petBD* (Figure 2E) and *ycf19* mentioned above and *cpcG1*, which are also preceded by RNase-E-dependent PSS. Hence, these 3’ UTRs are top candidates to act as 3’ end-derived sRNAs, analogous to several such riboregulators identified in enterobacteria (60–62). In cyanobacteria, only a single 3’ UTR-derived sRNA is known thus far, ApcZ (63). Hence, our here presented dataset provides a valuable resource for the functional characterization of further such sRNAs.

The six RNase E cleavage sites identified directly upstream of a start codon belong to the genes *psbA2* (*slr1311*, photosystem II D1 protein), *cpcB* (*sll1577*, phycocyanin beta subunit), *sigG* (*slr1545*, sigma factor), *slr0373* (hypothetical protein), *rcp1* (*slr0474*, two-component response regulator, interacting with cyanobacterial phytochrome Cph1) and *slr1563* (a fructosamine kinase family protein). Of these, the first four are the first gene within the respective transcriptional unit, whereas the latter two are the second or third and the identified cleavage sites are located exactly at the end of an intergenic region. Cutting a 5‘ leader can serve as a post-transcriptional regulatory mechanism by lowering translation initiation efficiency or by initiating further degradation, demonstrated in *Synechocystis* for the *psbA2* (21) and *psaL* mRNAs (28) and hence very likely also for the here detected additional genes. RNase-E-dependent cleavage sites within intergenic regions indicates a wider role of *Synechocystis* RNase E in the process of operon discoordination, as it was shown for the *rimO*-*crhR* dicistron (64).

### RNase E facilitates degradation of RNA fragments with structured 5’ ends

To identify factors determining the cleavage position of RNase E, we analysed PSS accumulating in *rne*(WT) or *rne*(Ts) both regarding their primary sequence as well as their calculated minimal folding energy (ΔG), which is indicative of the presence of structured regions (Figure 5). Analysing the minimal folding energy (ΔG) using a 25 nt sliding window approach revealed that the regions surrounding PSS accumulating in *rne*(WT) were less structured than for shuffled sequences or randomly picked positions (Figure 5A, Supplementary Figure 10). This is in concordance with the known preference of RNase E for single-stranded regions (9, 29). Regions exactly downstream of PSS with a higher read count in *rne*(Ts) are more structured than the average minimal folding energy and there is almost no sequence preference for the respective regions. The accumulation of these RNA fragments in *rne*(Ts) after RNase E inactivation indicates that these PSS are product of another ribonuclease, but that RNase E activity is necessary to further degrade them (Figure 6A).

**Figure 5.**
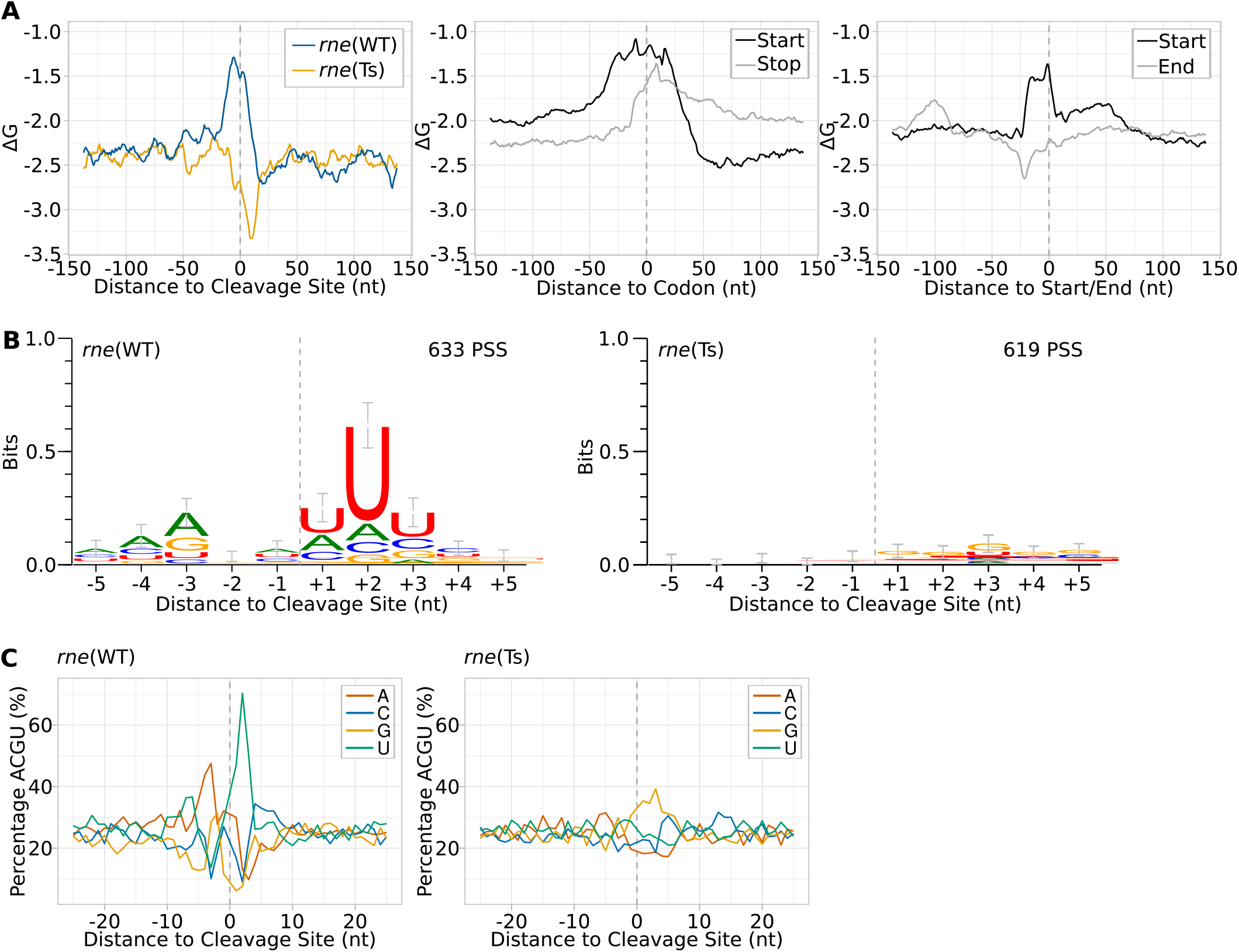
Inference of a sequence signature and folding potential around cleavage sites. (**A**) Analysis of minimal folding energy (ΔG) around detected cleavage sites, start and stop codons and annotated starts and ends of transcriptional units. Minimal folding energy was calculated at each nucleotide position using a sliding window of 25 nt and a 1 nt step size for a segment of 150 nt up- and downstream of the respective element. (**B**) Sequence logos of PSS accumulating in *rne*(WT) or *rne*(Ts) after heat treatment. Sequences were aligned according to the detected cleavage site. Error bars were calculated by the WebLogo tool and represent an approximate Bayesian 95% confidence interval. (**C**) Nucleotide composition around detected PSS.

**Figure 6.**
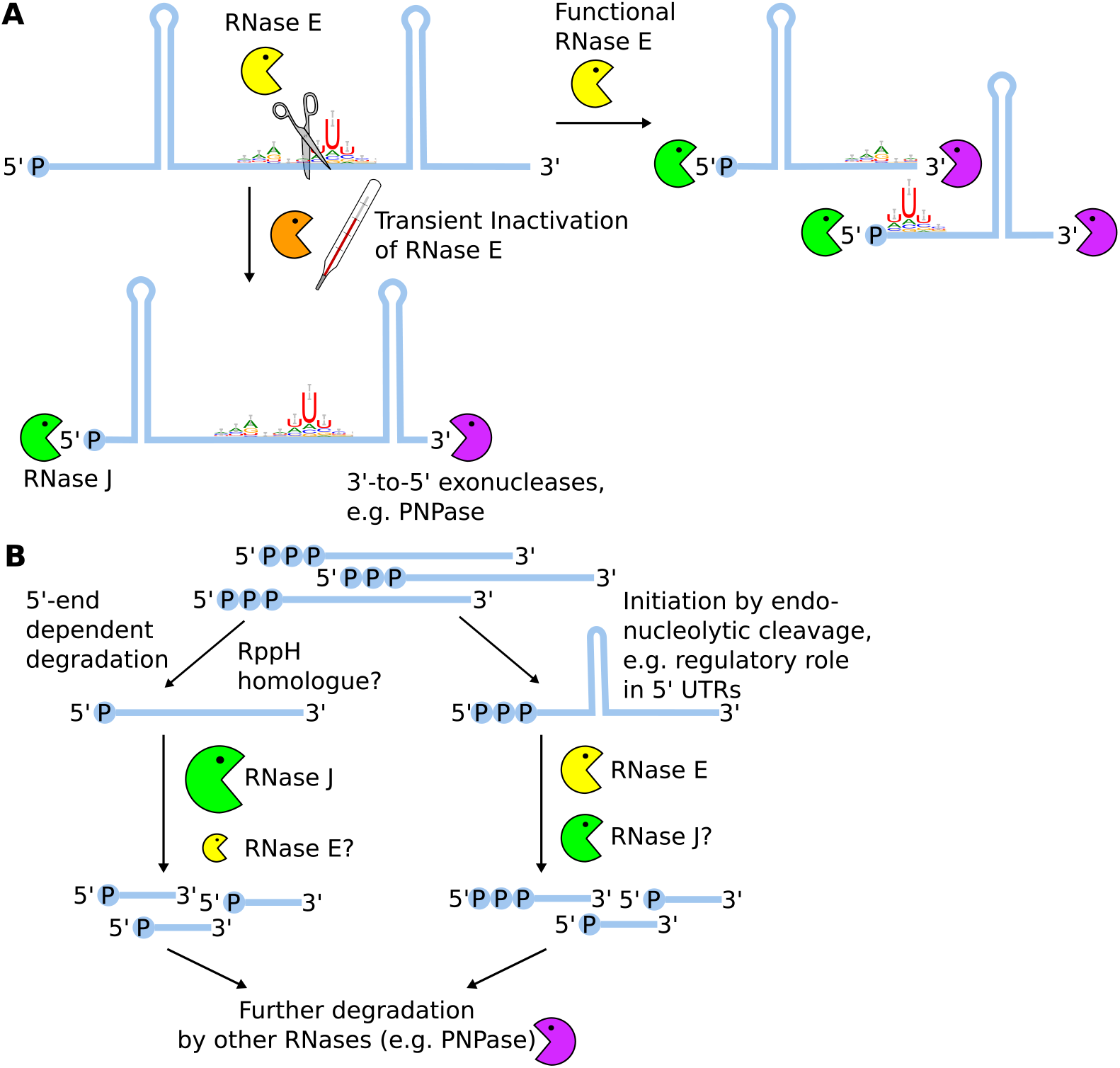
Hypothetical schemes for the coordinated action of RNase E and J in transcript turnover in *Synechocystis*. (**A**) RNase E activity is needed to make structured RNA species accessible for exonucleolytic activity of both 5‘-to-3‘ as well as 3‘-to-5‘ exonucleases. (**B**) Transient inactivation of RNase E did not impair 5‘-end dependent RNA degradation, hinting towards an important role of RNase J therein. However, RNase E inactivation strongly affected post-transcriptional regulation via regulatory RNA elements such as 5‘ UTRs.

### Less pronounced secondary structure around start and stop codons might attract RNase E cleavage

We analysed the minimal folding energy surrounding sequences start and stop codons as well as starts and ends of the TUs defined by Kopf et al. (46). We observed that regions around start and stop codons as well as TU starts are less structured than other sections in the genome (Figure 5A). Shortly before the ends of TUs, ΔG was slightly lowered, indicative of the presence of terminator structures. The lowered propensity for secondary structures around start and stop codons may contribute to the preference of RNase E for those positions.

### RNase-E-dependent PSS reveal a role of the A-U-clamp for positioning cleavage

Sites with higher PSS counts in *rne*(WT) were enriched for adenine residues at positions 4 nt and 3 nt upstream of the cleavage site (Figure 5B), while the two bases exactly upstream of the cleavage site showed no base preference. The three bases following the cleavage site were enriched for uracil residues, especially at position +2 relative to the cleavage site. These findings are also reflected by the overall nucleotide content of the neighbouring positions (Figure 5C). The created A-U-clamp appears to be a low-specificity and flexible way of positioning the actual cleavage site. Although we cannot discriminate between PSS resulting from endonucleolytic activity of RNase E and RNase J, which exhibited similar processing activities *in vitro* (20), such a sequence preference was not yet reported for RNase J. Thus, the identified A-U-clamp is likely RNase E specific and the +2 uridine ruler mechanism identified for *Salmonella* RNase E is conserved in cyanobacterial RNase E (8). However, enterobacterial and cyanobacterial RNase E differ in the nucleotide composition upstream of the cleavage site: *Salmonella* RNase E possesses a slight preference for a guanine residue at position -2 (8). The key importance of this guanine residue was shown for *E. coli* RNase E when acting on short RNA substrates lacking additional targeting factors such as secondary structures (65). When guanine was exchanged in these substrates for adenine, the cleavage rate of the assayed substrate was reduced by a factor of 12 (65). Another study found that the respective guanine residue is crucial for cleavage if no secondary structures are present in close distance which could guide RNase E to its cleavage site (13). According to these findings, enterobacterial and cyanobacterial RNase E both prefer cleavage after purine bases and rely, to a certain extent, on a uracil residue two nucleotide downstream of the cleavage site. Interestingly, the consensus motif for RNase-E-dependent cleavage sites determined for the GC-rich bacterium *Rhodobacter* differs strongly from the one presented here (50). A possible reason is that the positioning of RNase E cleavage sites in a GC-rich organism relies more strongly on alternative factors such as 5’ sensing or the presence of neighbouring secondary structures.

### RNase E activity is not rate-limiting for 5’-end dependent RNA degradation

Contrasting to what would be expected after the inactivation of a 5‘-P dependent RNase, the relative number of PSS reads compared to the number of TSS or transcript reads did not increase in *rne*(Ts) samples after the heat shock compared to the ratios in all other sequenced samples (Figure 2C). We assume that the transient inactivation of RNase E has no effect on the transformation of 5‘-PPP ends to 5‘-P ends, i.e. the action of a potential RNA pyrophosphohydrolase (RppH) homologue. Hence, the ratio of PSS reads to TSS reads at TSS positions is an indicator of the velocity and the extent of 5‘-end-dependent degradation. We did not observe a significant difference between PSS/TSS ratios at TSS positions between different strains at the used conditions.

If RNase E activity was rate-limiting for 5’-end dependent RNA degradation, the number of 5‘-P ends relative to 5‘-PPP ends or transcript reads should rise after RNase E inactivation. Such an effect was observed after deletion of RNase J1 and J2 in *Staphylococcus aureus* (66). This implies that RNase J might be the main ribonuclease performing 5‘-end dependent RNA degradation also in cyanobacteria (Figure 6B). However, a milder effect on the transcriptome was found for a knock-down of RNase J in *Synechocystis* than for RNase E (20). In light of this finding, we assume that both cyanobacterial RNase E and RNase J can partially substitute for each other and that their combined pool is sufficiently large to compensate for an inactivation of one of them, in regard to 5‘-end dependent degradation.

### Transient inactivation of RNase E leads to downregulation of *sigA*

Transcripts of several sigma factors, such as *sigA* and *sigE* are down-regulated in *rne*(Ts) compared to *rne*(WT) after heat treatment. The strong downregulation of *sigA* (log2FC=-2.0, p.adj=1.1*10^−41^), encoding the principal sigma factor SigA, indicates that RNase E inactivation confronts the cell with a strong stress in line with previous observations that it is downregulated in response to stress conditions such as heat, high salinity or photooxidative stress (reviewed in (67)). Interestingly, no other, alternative, sigma factor was significantly upregulated compared to the control strain. In contrast, growth at 39°C appeared to be only a minor stress for the congenic control strain, judged by the negligible effect on *sigA* transcription (log2FC=-0.6, p.adj=4.5*10^−4^) and moderate upregulation of heat-responsive sigma factor *sigB* (log2FC=1.2, p.adj=1.9*10^−19^). Interestingly, an asRNA complementary to the 3‘ region of *sigA* was strongly upregulated in *rne*(Ts) after the heat shock (TU3505, Supplementary Figure 11). This asRNA originates from a TSS leading to an overlap with the rRNA precursor. Also, several PSS detected in this asRNA were associated with divergent read counts between *rne*(WT) or *rne*(Ts), which is indicative of a direct action of RNase E on this transcript. Hence, this asRNA, which may impact the processing of the rRNA precursor and the level of the *sigA* mRNA level, is also a substrate for RNase E.

### Effect of RNase E inactivation on non-coding RNAs substantiate its role in post-transcriptional regulation

The role of RNase E in post-transcriptional regulation in enterobacteria is well established, while a few examples indicated a likely important function also in *Synechocystis* (21, 27, 28). Here, we noticed several non-coding RNAs were differentially expressed in the *rne*(WT) versus *rne*(Ts) comparison after one hour heat treatment, e.g. 16.0% of all annotated asRNAs and 16.4% of all annotated ncRNAs (Table 3). Hence, we decided to elucidate the impact of RNase E on these RNAs more closely.

A well-established target of *Synechocystis* RNase E is the 5‘ UTR of *psbA2*, encoding the photosystem II D2 protein, which is degraded by RNase E during the acclimation to darkness (20, 21, 29). *In vitro* cleavage assays conducted on a 35-mer identified major cleavage positions downstream of positions 27 and 31 (20, 29). Indeed, we detected PSS within the *psbA2* 5‘ UTR which were associated with divergent read counts between *rne*(WT) and *rne*(Ts), albeit at different positions than mapped *in vitro* (Figure 4B). We identified such a PSS at position 17 (pos. 7,197 on the chromosome).

PsrR1 is an sRNA upregulated in response to high light and CO_2_ depletion. PsrR1 regulates the expression of several photosynthesis-related proteins, e.g. PsaL, CpcA, PsbB and PsaJ on post-transcriptional level (28). For *psaL* mRNA, it was shown that this regulation involves RNase E. In our data set, PsrR1 was strongly downregulated after the heat shock. In *rne*(Ts), PsrR1 levels were further reduced than in *rne*(WT). PsrR1 was predicted to interact with *cpcBA* by base-pairing close to the start codon of both genes (28). Here, we observed PSS with enhanced read counts in *rne*(WT) compared to *rne*(Ts) after the heat shock which were located close to the start codons of *cpcA* and *cpcB* and thereby potentially removing the AUG from the respective coding region (Supplementary Figure 12). This implies that, similar to PsaL, the translation of CpcA and CpcB is also regulated by a combined action of PsrR1 and RNase E.

PmgR1 is an sRNA involved in the switch to photomixotrophic growth (68). PmgR1 was upregulated in WT, *rne*(WT) and *rne*(Ts) after the heat shock (Figure 4C, 4F). This upregulation was more strongly pronounced in *rne*(WT) than in *rne*(Ts). We identified several PSS in PmgR1 which accumulated in WT and *rne*(WT) compared to *rne*(Ts), indicative of a processing of PmgR1 by Rnase E. When analysed via northern blot hybridisation, a shorter version of PmgR1 produced in WT and *rne*(WT) was missing in *rne*(Ts) (Figure 4F). Direct RNA targets of PmgR1 are unknown and its mechanism remains to be elucidated. Processing by RNase E might yield mature, active PmgR1 or, contrary, inactivate the sRNA, e.g. by removing a putative seed region.

The transcription and translation of *gifB* encoding the glutamine synthetase inactivating protein factor IF17 are controlled by the transcription factor NtcA and a glutamine riboswitch, respectively (69, 70). The *gifB* transcript accumulates strongly in *rne*(Ts) after the heat shock, but not in *rne*(WT) or WT (Figure 4D, Figure 4G). Concurrently, several PSS with higher counts for *rne*(Ts) than *rne*(WT) after the heat shock were detected in the *gifB* CDS and riboswitch, and in the asRNA *sll1515-as2*, which overlaps the *gifB* 5’ UTR. The single PSS within the riboswitch is located at position +35, which was experimentally demonstrated to be essential for riboswitch function (69). The exact mechanism of action has remained unknown for this riboswitch. Based on the identified PSS in a functionally relevant domain, we hypothesize that this riboswitch might be dual-acting, reminiscent of the lysine sensing riboswitch *lysC* present in *E. coli*, which is both controlling translation of the downstream CDS as well as exposing RNase E cleavage sites upon lysine binding (71, 72). Additionally, RNase-E-dependent cleavage sites elsewhere within the mRNA and *sll1515-as* might affect *gifB* mRNA stability further.

The autoregulation of the *rne* transcript in *E. coli* is another example of 5’ UTR-mediated transcript level regulation (54). Using TIER-seq, we captured several RNase-E-dependent cleavage positions in the 5‘ UTR of *rne* (Figure 4E). Together with the enhanced accumulation of the *rne-rnhB* transcript upon RNase E inactivation (Figure 1E), this provides strong evidence for the autoregulation of *rne* levels by RNase E also in *Synechocystis*.

Additional genes whose transcript level seem to be regulated via an RNase E cleavage within their 5‘ UTR are *sll0547*, coding for an unknown protein and *rbp2* (*ssr1480*), encoding an RNA-binding protein involved in targeting mRNAs for photosynthetic proteins to the thylakoid membrane (73). Further examples for asRNAs in which RNase-E-dependent PSS were found are *slr0261-as*, which is located antisense to *ndhH*, encoding NADH dehydrogenase subunit 7, as well as the previously studied asRNAs IsrR (74) and PsbA3R (27).

## CONCLUSION

Our findings demonstrate that a substitution homologous to the *rne-3071* mutation of *E. coli* RNase E can be successfully introduced in a cyanobacterial RNase E, if combined with one of three detected spontaneous secondary mutations. This observation opens up an important avenue for the *in vivo* investigation of RNase E homologues that are more distantly related to the archetypical enterobacterial model. The generated strain enabled us to identify 1,472 RNase-E-dependent cleavage sites. Cleavage by an endoribonuclease results in a 5‘ and a 3‘ fragment, which may be rapidly further processed by other RNases. tagRNA-Seq only captures stable 5‘ fragments, but not 3‘ ends, which could be added in future studies by using an RNA-Seq approach also capturing 3‘ ends (75). Whereas cyanobacterial RNase E activity is not rate-limiting for 5‘-end dependent RNA degradation, we showed it to be important for the degradation of RNA fragments with strong 5‘ secondary structures, which hints towards one possible reason why the enzyme is essential in *Synechocystis*. Our data show that cyanobacterial RNase E differs from enterobacterial RNase E slightly in its target affinity, by preferring adenine upstream of cleavage sites instead of guanine. Future work has to elucidate if other targeting mechanisms known for enterobacterial RNase E, e.g. 5’ sensing and 5’ bypass, are present in the compact cyanobacterial RNase E and which relevance they might have. The created mutant strain, *rne*(Ts), represents a promising tool for the future analysis of post-transcriptional regulation as well as the maturation and action of regulatory ncRNAs. Exploring the impact of transient RNase E inactivation under different environmental conditions will help to further understand the function of this widely conserved and versatile endoribonuclease in cyanobacteria.

## Supporting information

.pdf file with Supplemental Methods, Results, Tables S1, S2 and Figures

.xlsx file with Tables S3 - S12

.xlsx file with Tables S13 - S14

.xlsx file with Tables S15 - S17

.xlsx file with Tables S18 - S25

.gff file with coordinates of TSS and PSS

## DATA AVAILABILITY

Galaxy workflows used for raw read processing and several downstream analysis steps are available (https://usegalaxy.eu/u/ute-hoffmann/w/tier-seqprocessing1, https://usegalaxy.eu/u/ute-hoffmann/w/tier-seqtranscript, https://usegalaxy.eu/u/ute-hoffmann/w/tier-seqpss-tss and https://usegalaxy.eu/u/ute-hoffmann/w/tiertranscript-coverage).

All further code used for data processing and analysis are available in the GitHub repository (https://github.com/ute-hoffmann/TIER-synechocystis).

## ACCESSION NUMBERS

FastQ files from RNA sequencing have been deposited with the Gene Expression Omnibus (GEO) Data bank under accession number GSE180316.

## SUPPLEMENTARY DATA

Supplementary Data are available at NAR online.

Supplement.pdf - .pdf file with Supplemental Methods, Results, Tables S1, S2 and Figures

SuppTables_1_S3-S12.xlsx - .xlsx file with Tables S3 – S12

SuppTables_2_S13-S14.xlsx - .xlsx file with Tables S13 – S14

SuppTables_3_S15-S17.xlsx - .xlsx file with Tables S15 – S17

SuppTables_4_S18-S25.xlsx - .xlsx file with Tables S18 – S25

TSS_PSS_rneAnalysis.gff - .gff file with coordinates of TSS and PSS

## FUNDING

This work was supported by the Deutsche Forschungsgemeinschaft Research Training Group MeInBio [322977937/GRK2344] to AW, WRH and RB and by Deutsche Forschungsgemeinschaft Grant STE 1192/4-2 to CS.

## CONFLICT OF INTEREST

The authors declare that there is no conflict of interest.

## ACKNOWLEDGEMENT

We thank J. Behler for providing the plasmid for the chromosomal integration of a 3xFLAG tag N-terminally of *rne*. We thank W. Bigott and S. Oeser for technical assistance, S. Oeser and N. Schuergers for fruitful discussions and M. Treppner for biostatistical advice.

The authors acknowledge the support of the Freiburg Galaxy Team: Björn Grüning, Bioinformatics, University of Freiburg (Germany) funded by the Collaborative Research Centre 992 Medical Epigenetics (Deutsche Forschungsgemeinschaft [SFB 992/1 2012]) and Bundesministerium für Bildung und Forschung BMBF grant 031 A538A de.NBI-RBC.

## Notes

https://www.ncbi.nlm.nih.gov/geo/query/acc.cgi?acc=GSE180316

### Competing Interest Statement

The authors have declared no competing interest.

