## Supplementary material for "Transcriptome-wide *in vivo* mapping of cleavage sites for the compact cyanobacterial ribonuclease E reveals insights into its function and substrate recognition": .pdf file with Supplemental Methods, Results, Tables S1, S2 and Figures

**Supplemental Information**

**Supplemental Methods**

**Supplemental Results**

**Supplemental Tables**

**Supplemental Figures**

**Supplemental References**

### SUPPLEMENTAL METHODS

#### 1 Detailed description of preliminary read processing

RNA-seq read quality was checked in the beginning and after each step using FastQC (http://www.bioinformatics.babraham.ac.uk/projects/fastqc). Reports were summarised with MultiQC (1). To separate PSS, TSS and transcript pattern reads, BBDuk was used (2) (https://www.sourceforge.net/projects/bbmap/). After upload to the Galaxy web platform (3), files were decompressed and read identifiers were modified using awk (Galaxy v1.1.2, https://github.com/bgruening/galaxytools) to be compatible with UMI-tools deduplicate with the following expression:

*{if($0 ~ /^@/) {val=$0; split(val,vector," "); split(vector[1], tag, ":"); print vector[1] "_" tag[8]} else {print $0} }*

Resulting files were further processed using a workflow (https://usegalaxy.eu/u/ute-hoffmann/w/tier-seqprocessing1) encompassing adapter trimming using Cutadapt (4). Trimmed reads were mapped to the *Synechocystis* genome with Bowtie2 and very-sensitive settings (5–7). PCR duplicates were removed using UMI-tools deduplicate with parameters (--method directional --edit-distance-threshold 1 --soft-clip-threshold 4 --subset 1.0) (8). Samtools stats and bamtools filter were run for control and filtering of the resulting .bam files (9–11). Reads with a mapping quality above 20 were sorted according to data type (TSS, PSS, unspecified transcript reads) and used as input for further workflows ( https://usegalaxy.eu/u/ute-hoffmann/w/tier-seqtranscript, https://usegalaxy.eu/u/ute-hoffmann/w/tier-seqpss-tss and https://usegalaxy.eu/u/ute-hoffmann/w/tiertranscript-coverage). In *Synechocystis*, two almost identical rRNA loci exist. The same holds for the coding regions of genes *psbA2* and *psbA3*, encoding photosystem II D1 proteins. When analysing the respective regions, multi-mapped reads were included in downstream analyses by additionally considering reads with a mapping quality of exactly one.

#### 2 Additional description of classification of TSS and PSS and downstream analyses

htseq-count files of transcript data were analysed using DESeq2 (v1.30.1) (12) (|log2FC| > 0.8 and p.adj < 0.05). Functional enrichment and gene set enrichment analyses (GSEA) were performed using clusterProfiler (v3.18.1) (13, 14) based on KEGG (15, 16) and Gene Ontology (GO) (17, 18) annotation. The latter was obtained from CyanoBase (19) and UniProt (20).

Since the enzymatic steps of adapter ligation and the subsequent removal of unligated 5‘-P RNA species are not 100% efficient, a certain amount of PSS RNA ends are ligated to TSS tags during tagRNA-Seq library preparation. TSS RNA ends may be converted to 5‘-P ends in living cells or during RNA extraction and library preparation. Hence, true TSS ends are present in the PSS data sets and vice versa. To classify TSS and PSS, the bedtools genome coverage 5’ ends files were downloaded and positions at which a low number of reads mapped were filtered out using edgeR (v3.32.1) function *filterByExpr()* with default settings (21). We used DESeq2 to distinguish between PSS and TSS positions to take advantage of the model underlying DESeq2 and therefore be able to assign probability values to the categorisation of a certain position as PSS or TSS. Size factors of PSS and TSS data sets were determined for the two data set types separately using DESeq2. These size factors were used in a subsequent DESeq2 analysis combining both data sets to test which genomic positions were significantly enriched in the PSS or the TSS data sets. Cut-off values of |log2FC| > 0.8 and p.adj < 0.05 were used for TSS/PSS classification (Supplementary Figure 5A).

We observed a high amount of false positive TSS positions in highly transcribed genomic regions. In addition, some potential TSS positions were located at genomic regions without detectable transcription, which we also assume to be false positives. To control for false positive TSS positions, edgeR *filterByExpr()* was used to filter out lowly populated genomic positions in transcript coverage files. This was followed by normalization of the transcript coverage and the TSS 5' ends data using DESeq2. The mean read count at each genomic position of all samples were calculated for transcript coverage and TSS 5' end counts separately. Only positions at which the ratio of the mean of TSS 5' ends divided by the mean of transcript coverage was above 0.02 and below 2.00 and which were located within 20 nt of a TSS annotated by Kopf et al. (22) were used in downstream analyses. These thresholds were determined by examination of the *psbA2* TSS region and comparison to experimentally validated TSS (23). PSS might also be affected by similar effects. However, since we were interested in degradation intermediates and these are often lowly abundant, we decided to not apply similar cut-offs and follow, in respect to PSS, an inclusive strategy.

Several Bioconductor (24) packages were used in downstream analyses, namely tidyverse (v1.3.0) (25), genomicRanges (v1.42.0) (26) and rtracklayer (v1.50.0) (27). For visualisation, data was averaged among triplicates, prepared using an R script and visualised with Artemis (28–31). Figures were created and analysed using Fiji (v2.0.0-rc-69/1.52p) (32) and Inkscape (v1.0.2).

### SUPPLEMENTAL RESULTS

#### 1 Enrichment analysis of newly identified PSS

The set of coding regions overlapping with the newly identified PSS were enriched for KEGG pathways and GO terms related to photosynthesis and the ribosome (Supplementary Figure 5C, Supplementary Tables S3, S4). Those terms include highly abundant proteins in cyanobacteria. When testing if a certain set of genes is enriched among the most highly transcribed genes in our data set according to the average base mean divided by the features' length, the KEGG term "photosynthesis" was enriched (p.adj = $1.7 * 10^{-3}$) and GO terms "photosynthesis" (p.adj = $8.5 * 10^{-5}$), "thylakoid membrane" (p.adj = $1.2 * 10^{-2}$) and "protein-chromophore linkage" (p.adj = $4.9 * 10^{-2}$) (Supplementary Tables S5, S6). In the described data set, the most highly expressed genes encoded phycobilisome or photosystems 1 and 2 subunits. This matches the enriched terms for the set of coding sequences in which PSS are located. Based on this finding, we assume that processing sites in lowly abundant transcripts were possibly partially not captured using tagRNA-Seq.

#### 2 Enrichment analysis of transcriptomic changes before and after heat shock

To further analyse the transcriptional changes after the heat shock, we performed functional enrichment analyses and gene set enrichment analyses (GSEA) with Gene Ontology (GO) and KEGG terms (Supplementary Figure 6, Supplementary Tables S7 – 10; p.adj < 0.05). In both strains, genes related to the KEGG terms "Oxidative phosphorylation" (syn00190; *rne*(WT): p.adj = $5.0*10^{-7}$, *rne*(Ts): p.adj = $1.8*10^{-8}$)

were depleted after the heat shock. This is complemented by a multitude of GO terms affected in both strains by the heat treatment. Several further KEGG terms are depleted in *rne*(Ts) after the 39°C incubation.

### 3 Analysis of RNase-E-dependent PSS within rRNA loci

RNase E is known to be involved in 5S, 16S and 23S rRNA maturation in other bacteria, for instance *E. coli* and *M. smegmatis* (33–35). Only few PSS detected within the rRNA precursor transcript accumulated in *rne*(Ts) or *rne*(WT) after the heat shock. Major maturation intermediates were not affected. However, a PSS potentially corresponding to a lowly abundant maturation intermediate of 5S rRNA was enriched in *rne*(WT) compared to *rne*(Ts). Furthermore, this was also the case for several PSS which were detected within 23S rRNA. It has to be noted that rRNA was depleted prior to library preparation, which is reflected by relatively low read numbers for 16S rRNA and 23S rRNA. Additionally, RNA-seq data was prepared from samples after heat treatment for only one hour, which might not be a sufficient long period of time to observe a significant accumulation of respective precursors.

### 4 Influence of RNase E inactivation on specific biological processes

To establish the impact of RNase E inactivation on specific biological processes, we performed GSEAs to identify pathways which might be regulated by RNase E (Supplementary Figure 8B, Supplementary Tables S20 – S23, p.adj < 0.05). As a complementary analysis, we performed functional enrichment analyses of the sets of coding sequences overlapped by PSS (Supplementary Figure 8C, Supplementary Tables S24, S25). On transcript level, only few GO terms and no KEGG terms were differentially enriched in both strains before the heat shock, whereas many GO and KEGG terms differed significantly between *rne*(WT) and *rne*(Ts) after the heat shock. This corroborates further that the introduced point mutations did not affect the transcriptome strongly at standard growth temperature. After the heat shock, photosynthesis-associated KEGG-terms (“Photosynthesis”: syn00195; p.adj = $1.2 * 10^{-6}$, “Photosynthesis – antenna proteins”: syn00196; p.adj = $3.5 * 10^{-3}$) were enriched on a transcript level in *rne*(WT), whereas being overrepresented among coding sequences overlapping with PSS accumulating in *rne*(Ts) (“Photosynthesis”: syn00195; p.adj = $1.0 * 10^{-3}$, “Photosynthesis – antenna proteins”: syn00196; p.adj = $1.8 * 10^{-2}$). Additional to related GO terms, the term “Nucleic acid binding” (p.adj = $4.0 * 10^{-2}$) was found to be enriched in coding sequences overlapping with *rne*(Ts) PSS. This term comprises, among others, the following genes: *slr0083* (*crhR*, RNA helicase), *ssr1480* (*rbp2*) and *sll0517* (*rbp1*).

**SUPPLEMENTAL TABLES**

**Table S1.** Oligonucleotides used in this study.

| Name | Sequence | Used for |
|---|---|---|
| P01 | GGCGTATCACGAGGCCCTTTCGTCTATCACCCTCAATGAACAGC | Amplification of flank 1 for homologous recombination of N-terminally 3xFLAG-tagged RNase E |
| P02 | TGCCGCTGGTGAGAACTGCCTAAACTGCCCG | Amplification of flank 1 for homologous recombination of N-terminally 3xFLAG-tagged RNase E |
| P03 | GAATATATTTATGGATTATAAAGATCATGATGGCGATTATAAAGATCATGATATTGATTATAAAGATGATGATGATAAACCAAAACAAATTGTCATTGCTG | Amplification of flank 2 for homologous recombination of N-terminally 3xFLAG-tagged RNase E |
| P04 | GGCCTTTTTACGGTTCCTGGCCTTTCCCCGCCAATATTTCTCAG | Amplification of flank 2 for homologous recombination of N-terminally 3xFLAG-tagged RNase E |
| P05 | AGTTTAGGCAGTTCTCACCAGCGGCAACCGC | Amplification of kanamycin resistance cassette for N-terminally 3xFLAG-tagged RNase E |
| P06 | TTATTCAAAGCCGGCCGCCGTCCCGTCAAGT | Amplification of kanamycin resistance cassette for N-terminally 3xFLAG-tagged RNase E |
| P07 | ACGGGACGGCGGCCGGCTTTGAATAAGGTAG | Amplification of *slr1129* promoter region |
| P08 | CCATCATGATCTTTATAATCCATAAATATATTCCTCAAAAGGC | Amplification of *slr1129* promoter region |
| P09 | CGTGTCGACGCAGTTAGCTTGGGGCTGG | Amplification of *rne-rnhB* locus with XhoI and SalI restriction sites |
| P10 | GCTCTCGAGGAAACGGCGGCTGAATG | Amplification of *rne-rnhB* locus with XhoI and SalI restriction sites and testing for full seggregation |
| P11 | CGAAAAAAATTCCTTTATCCACGTCAG | Primer for mutagenesis G63S |
| P12 | GTGTCCCCAATGTTAATAAAAG | Primer for mutagenesis G63S |
| P13 | AAATGGCTTTTTTCACGTCAGTGAC | Primer for mutagenesis I65F |
| P14 | TTTTCGGTGTCCCCAATG | Primer for mutagenesis I65F |
| P15 | TTTTCACGTCAGTGACCTCGGC | Primer for mutagenesis double mutation G63S I65F |
| P16 | AAGGAATTTTTTTCGGTGTCCCCAATG | Primer for mutagenesis double mutation G63S I65F |
| P17 | TATCCGGGAAGCCCTTAAAC | Testing insertion direction and sequencing |
| P18 | GTTCATCATGCCGTCTGTGATG | Testing insertion direction and sequencing |
| P19 | AGGGGTTACAGTCAGCTTGC | Testing insertion direction |
| P20 | CGAGGCCCTTTCGTCTGGAAGTTAACTATTCGTT | Amplification of upstream flank for homologous recombination for AQUA cloning |
| P21 | CGTTGAATATGGCTCATCCTAATACCCAAGGAATT | Amplification of upstream flank for homologous recombination for AQUA cloning |

| Name | Sequence | Used for |
|---|---|---|
| P22 | AATTCCTTGGGTATTAGGATGAGCCATATTCAACG | Amplification of kanamycin resistance cassette for AQUA cloning |
| P23 | GGTCGGTGTAGAGGAGTCGTTAGAAAAACTCATCG | Amplification of kanamycin resistance cassette for AQUA cloning |
| P24 | CGATGAGTTTTTCTAACGACTCCTCTACACCGACC | Amplification of downstream flank for homologous recombination for AQUA cloning |
| P25 | TTACGGTTCCTGGCCTTTCTGAGCATGGAAGTGGTGTC | Amplification of downstream flank for homologous recombination for AQUA cloning |
| P26 | GACACCACTTCCATGCTCAGAAAGGCCAGGAACCGTAA | Amplification of plasmid backbone for homologous recombination for AQUA cloning |
| P27 | AACGAATAGTTAACTTCCAGACGAAAGGGCCTCG | Amplification of plasmid backbone for homologous recombination for AQUA cloning |
| P28 | GTTGCAATTCCTTTTGGCCCAG | Test for full seggregation |
| P29 | AGTCCCAACTCCGTCAACTG | Sequencing |
| P30 | CCCAGTGATGGGTTGACAAT | Sequencing |
| P31 | AGCCTTGGGAAAACGGTACT | Sequencing |
| P32 | AGTCCCATGCCCGGTGGC | Sequencing |
| P33 | GGGCGTTATATGGTGTTG | Sequencing |
| H01 | TAATACGACTCACTATAGGGCAAGGTGGCCTACAGAA | Amplification of DNA template for *in vitro* transcription of probe complementary to 5‘ UTR of *rne* |
| H02 | GTAGTGAATCGCCGTAAGCAGCC | Amplification of DNA template for *in vitro* transcription of probe complementary to 5‘ UTR of *rne* |
| H03 | TAA TAC GAC TCA CTA TAG GGT AAT AGT AATGAC AGG CAG | Amplification of DNA template for *in vitro* transcription of probe complementary to CRISPR3 Spacers 1 – 4 |
| H04 | CTT TAG GTG GGC GTT GAC CT | Amplification of DNA template for *in vitro* transcription of probe complementary to CRISPR3 Spacers 1 – 4 |
| H05 | TAATACGACTCACTATAGGGGCACTGTCCTCACGCTCGC | Amplification of DNA template for *in vitro* transcription of probe complementary to RnpB |
| H06 | GAGTTAGGGAGGGAGTTGCGG | Amplification of DNA template for *in vitro* transcription of probe complementary to RnpB |
| H07 | TAATACGACTCACTATAGGGAAATCACTTCCAAACAACACCC | Amplification of DNA template for *in vitro* transcription of probe complementary to PmgR1 (Ncr0700) |
| H08 | GTACATTGAATACATGGAGCCGAAG | Amplification of DNA template for *in vitro* transcription of probe complementary to PmgR1 (Ncr0700) |
| H09 | TAATACGACTCACTATAGGGCGACTAGTTAGTAGGTAAG | Amplification of DNA template for *in vitro* transcription of probe complementary to *gifB* |
| H10 | GGATACAGAAAGTAAATCGTTC | Amplification of DNA template for *in vitro* transcription of probe complementary to *gifB* |

**Table S2.** Strains used in this study.

| Name | Description |
|---|---|
| 3xFLAG-*rne* | *slr1128*::*kanR 3xFLAG-rne*$^+$ |
| *rne*(WT) | Δ(*rne-rnhB*) *rnep*::*kanR* pVZ321Δ(9220-1203) (9219)::(*rnep-3xFLAG-rne*$^+$*-rnhB-rnhBt*) |
| *rne*(Ts) | Δ(*rne-rnhB*) *rnep*::*kanR* pVZ321Δ(9220-1203) (9219)::(*rnep*-3xFLAG-*rne*(Ts)-*rnhB-rnhBt*) |
| pVZΔKmR | *slr1128*::*kanR 3xFLAG-rne*$^+$ pVZ321Δ(9220-1203) |

**Overview Further Supplemental Tables**

For Tables S3 – S25, see separate Excel work books SuppTables_1_S3-S12.xlsx, SuppTables_2_S13-S14.xlsx, SuppTables_3_S15-S17.xlsx and SuppTables_4_S18-S25.xlsx.

**Supplementary Table S3.** Gene Ontology Functional Enrichment Analysis of coding features overlapped by PSS.

**Supplementary Table S4.** KEGG Functional Enrichment Analysis of coding features overlapped by PSS.

**Supplementary Table S5.** Gene Ontology Gene Set Enrichment Analysis analysing enrichment of terms among highly transcribed genes (baseMean/width).

**Supplementary Table S6.** KEGG Gene Set Enrichment Analysis analysing enrichment of terms among highly transcribed genes (baseMean/width).

**Supplementary Table S7.** Gene Ontology Gene Set Enrichment Analysis comparing *rne*(WT) before and after heat shock, higher values are enriched after 1h.

**Supplementary Table S8.** KEGG Gene Set Enrichment Analysis comparing *rne*(WT) before and after heat shock, higher values are enriched after 1h.

**Supplementary Table S9.** Gene Ontology Gene Set Enrichment Analysis comparing *rne*(Ts) before and after heat shock, higher values are enriched after 1h.

**Supplementary Table S10.** KEGG Gene Set Enrichment Analysis comparing *rne*(Ts) before and after heat shock, higher values are enriched after 1h.

**Supplementary Table S11.** DESeq2 analysis on level of RNA features comparing *rne*(WT) before and after heat shock (1h / 0h). RNA features are sorted according to increasing adjusted p values.

**Supplementary Table S12.** DESeq2 analysis on level of RNA features comparing *rne*(Ts) before and after heat shock (1h / 0h). RNA features are sorted according to increasing adjusted p values.

**Supplementary Table S13.** DESeq2 analysis on level of RNA features comparing *rne*(WT) and *rne*(Ts) at time point 0h. RNA features are sorted according to increasing adjusted p values.

**Supplementary Table S14.** DESeq2 analysis on level of RNA features comparing *rne*(WT) and *rne*(Ts) at time point 1h. RNA features are sorted according to increasing adjusted p values.

**Supplementary Table S15.** DESeq2 analysis on level of transcriptional units comparing *rne*(WT) and *rne*(Ts) at time point 0h. Transcriptional units are sorted according to increasing adjusted p values.

**Supplementary Table S16.** DESeq2 analysis on level of transcriptional units comparing *rne*(WT) and *rne*(Ts) at time point 1h. Transcriptional units are sorted according to increasing adjusted p values.

**Supplementary Table S17.** DESeq2 analysis for PSS comparing *rne*(WT) and *rne*(Ts) at time point 0h. PSS are sorted according to increasing adjusted p values.

**Supplementary Table S18.** DESeq2 analysis for PSS comparing *rne*(WT) and *rne*(Ts) at time point 1h. PSS are sorted according to increasing adjusted p values.

**Supplementary Table S19.** DESeq2 analysis for PSS comparing *rne*(WT) before and after heat shock (1h / 0h). PSS are sorted according to increasing adjusted p values.

**Supplementary Table S20.** Gene Set Enrichment Analysis analysing enrichment of certain type of RNA features comparing *rne*(WT) and *rne*(Ts) at time point 1h, higher values are enriched in *rne*(WT).

**Supplementary Table S21.** Gene Ontology Gene Set Enrichment Analysis comparing *rne*(WT) and *rne*(Ts) at time point 0h, higher values are enriched in *rne*(WT).

**Supplementary Table S22.** Gene Ontology Gene Set Enrichment Analysis comparing *rne*(WT) and *rne*(Ts) at time point 1h, higher values are enriched in *rne*(WT).

**Supplementary Table S23.** KEGG Gene Set Enrichment Analysis comparing *rne*(WT) and *rne*(Ts) at time point 1h, higher values are enriched in *rne*(WT).

**Supplementary Table S24.** Gene Ontology Functional Enrichment Analysis of coding features overlapped by PSS which are upregulated in *rne*(Ts) after 1h heat, universe: coding sequences generally overlapped by PSS.

**Supplementary Table S25.** KEGG Functional Enrichment Analysis of coding features overlapped by PSS which are upregulated in *rne*(Ts) after 1h heat, universe: coding sequences generally overlapped by PSS.

### SUPPLEMENTAL FIGURES

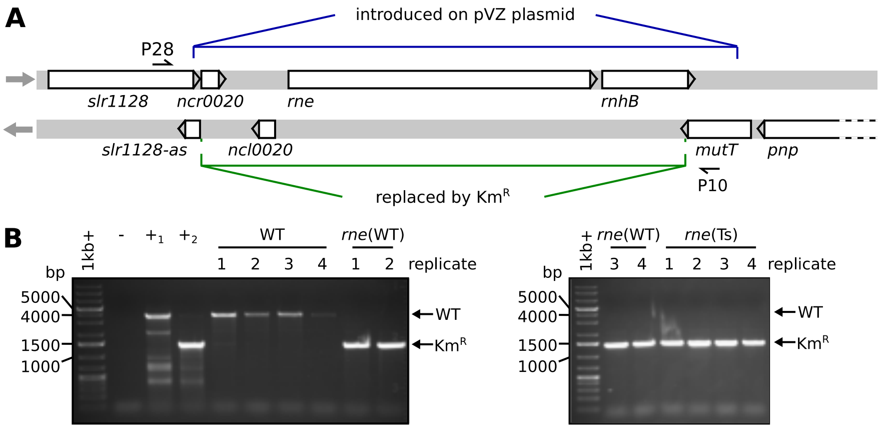


**Supplementary Figure 1.** Overview and verification of creation of mutant strains. (**A**) Overview of the genomic locus of *rne* and *rnhB*. Sites at which primer pair P10/P28 binds, which was used for verification of segregation, are indicated by small arrows. Regions which were introduced on conjugative plasmid pVZ321 and which were replaced by a kanamycin resistance cassette (Km$^{\mathrm{R}}$) by homologous recombination are indicated in blue and green, respectively. (**B**) Agarose TAE gels with products of PCR performed with primers P28/P10 to verify full segregation of cultures used for TIER-seq experiment. (-): Negative control, water was added instead of template, $(+_1)$: genomic wild-type DNA, $(+_2)$: genomic DNA of *rne*(WT). PCR products from the wild-type locus have a size of 3813 bp. The expected PCR product size is 1440 bp after introduction of the kanamycin resistance cassette. Arrows indicate the respective product sizes.

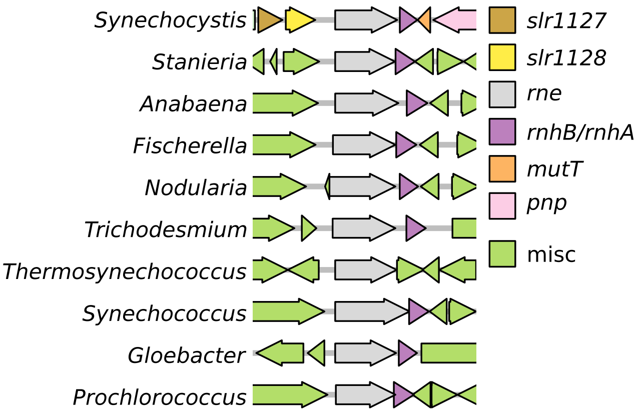


**Supplementary Figure 2.** Synteny analysis of *rne* and *rnhB*. Only genes present in proximity of *rne* in *Synechocystis* are labelled by name. All *rnhB*/*rnhA* homologues are annotated as *rnhB* except *Gloebacter rnhA*. The following species are shown in the Figure: *Synechocystis* sp. PCC 6803 (*Synechocystis*), *Stanieria cyanosphaera* PCC 7437 (*Stanieria*), *Anabaena* sp. PCC 7120 (*Anabaena*), *Fischerella* sp. NIES 3754 (*Fischerella*), *Nodularia spumigena* CCY9414 (*Nodularia*), *Trichodesmium erythraeum* IMS101 (*Trichodesmium*), *Thermosynechococcus elongatus* BP-1 (*Thermosynechococcus*), *Synechococcus elongatus* PCC 7942 (*Synechococcus*), *Gloeobacter violaceus* PCC 7412 (*Gloeobacter*), *Prochlorococcus marinus* str. MIT 9313 (*Prochlorococcus*). The following accessions from NCBI GenBank (36, 37) were used for analysis: BA000022.2 (5), CP003653.1, BA000019.2 (38), AP017305.1 (39), CP007203.2 (40), CP000393.1, BA000039.2 (41), CP000100.1, BA000045.2 (42), BX548175.1 (43).

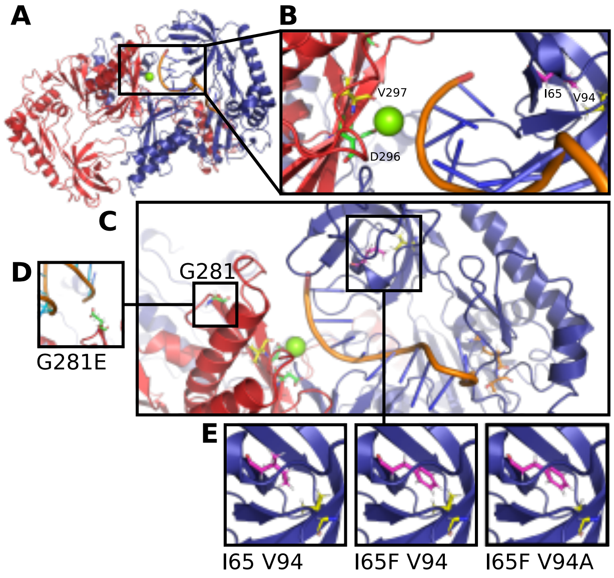


**Supplementary Figure 3.** Homology model of a dimer of *Synechocystis* RNase E, created using iTasser (44–46) and Pymol. Only the N-terminal protein part up to amino acid V496 is depicted since the last 178 amino acids of the protein show a very high flexibility. Protein crystal structures of *E. coli* RNase E were used to orient two monomers (PDB (47) ID: 6G63 (48), 2C0B (49)) and the RNA moieties co-crystallized with these structures were added (RNA moiety from 2C0B in all subfigures, RNA moiety from 6G63 for (D)). Green: Magnesium ion. (**A**) Overview of the dimer with two monomers depicted in red and blue. (**B**) RNA binding channel. (**C**) RNA binding channel in connection with the 5'-sensing pocket. (**D**) Localisation of compensatory mutation G281E. (**E**) Localisation of mutation I65F and compensatory mutation V94A.

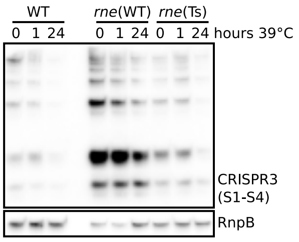


**Supplementary Figure 4.** Northern blot analysis of accumulation of mature CRISPR3 crRNAs, representative of 3 replicates.

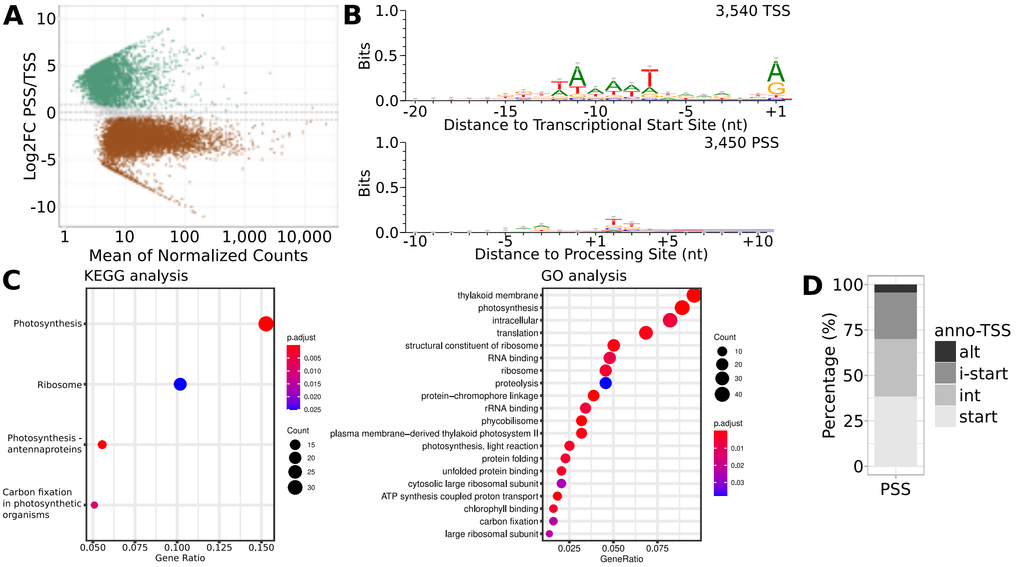


**Supplementary Figure 5.** Figures associated with deterimation of PSS and further characterization thereof. (**A**) MAplot of TSS and PSS peaks. (**B**) Sequence logo created from sequences surrounding all PSS and all TSS identified by tagRNA-Seq. Sequences were aligned according to the positioning of PSS or TSS. Error bars are automatically calculated by the WebLogo tool and represent an approximate Bayesian 95% confidence interval. (**C**) Functional enrichment analysis of coding sequences overlapped by all identified PSS using KEGG or GO terms. (**D**) Overlap of PSS identified by tagRNA-Seq with TSS annotated by Kopf et al. (22) (anno-TSS).

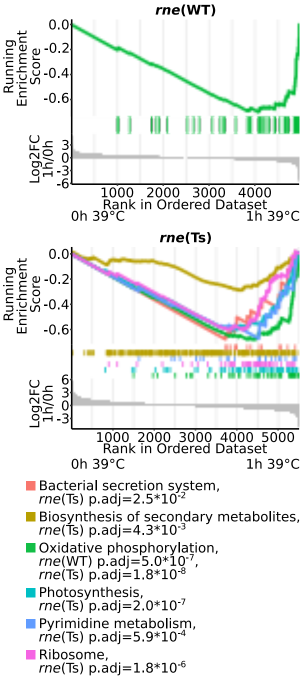


**Supplementary Figure 6.** Gene set enrichment analyses using KEGG terms comparing transcript composition before and after the heat shock for *rne*(WT) (upper panel) and *rne*(Ts) (lower panel).

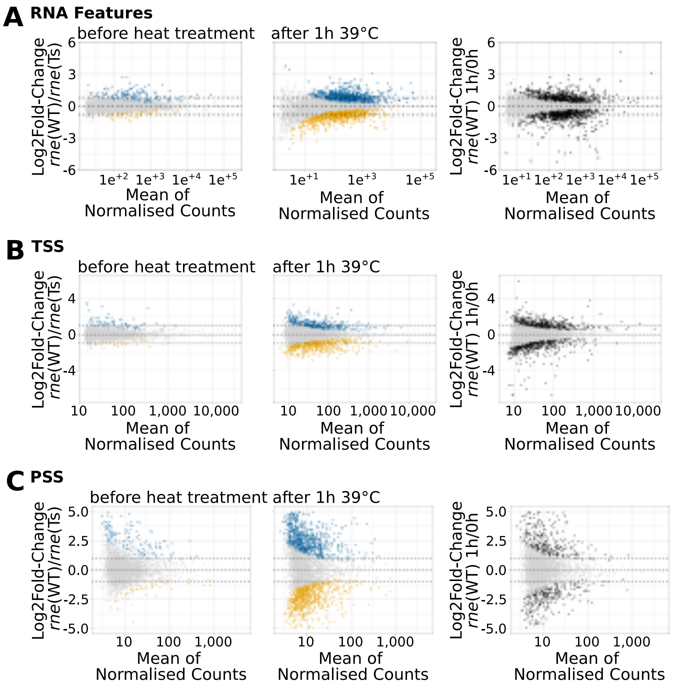


**Supplementary Figure 7.** MA plots of the TIER-Seq analysis. (**A**) Transcript data analysed on level of RNA features. (**B**) TSS. (**C**) PSS. Dotted lines indicate the chosen log2FC cut-off values (0.8 for transcript data, 1.0 for TSS and PSS). When points are coloured in blue, yellow or black, their associated adjusted p value is smaller than 0.05.

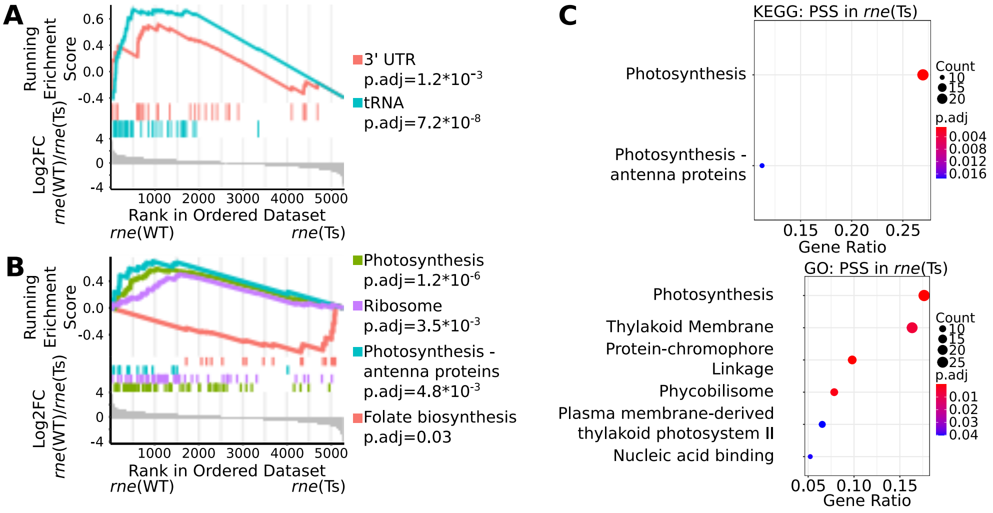


**Supplementary Figure 8.** Functional enrichment analyses comparing *rne*(WT) and *rne*(Ts) after one hour of heat treatment. (**A**) Gene set enrichment analyses (GSEA) testing distribution of different transcript types and RNA regions. (**B**) GSEA testing distribution of transcripts associated with a certain KEGG term. (**C**) Functional enrichment analysis of set of coding sequences overlapped by PSS accumulating in *rne*(Ts).

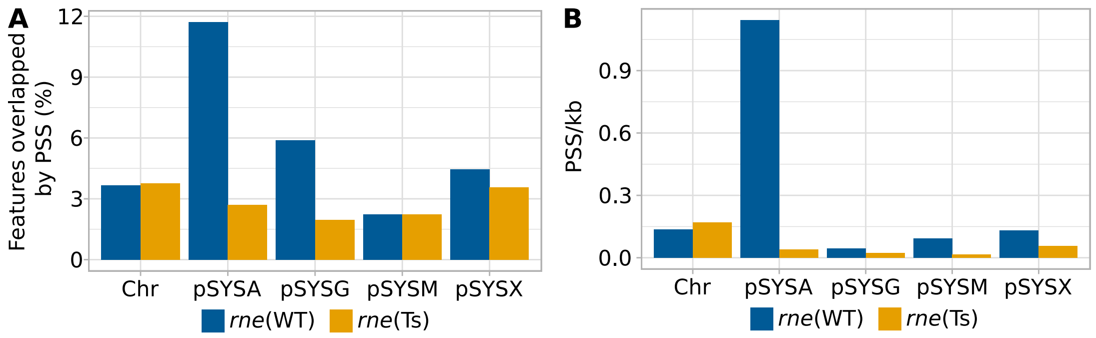


**Supplementary Figure 9.** Analysis of PSS overlapping different replicons. (**A**) Percentage of annotated features on different parts of genome overlapped by PSS comparing *rne*(WT) and *rne*(Ts) after the heat treatment. Note that CRISPR repeat and spacer sequences of each of the three CRISPR arrays were summarized as only one feature, respectively. (**B**) PSS per kilobase for different parts of genome comparing *rne*(WT) and *rne*(Ts) after the heat treatment. Chr: chromosome.

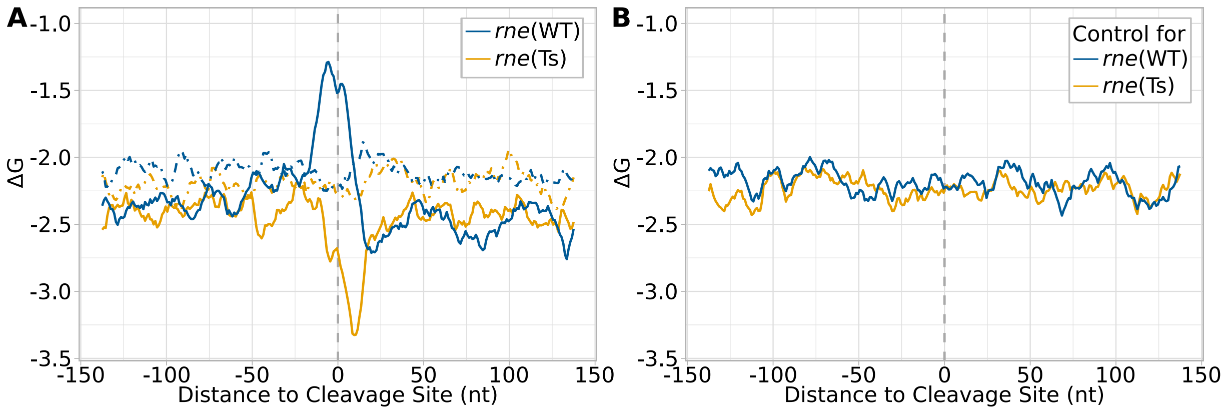


**Supplementary Figure 10.** Controls for minimal folding energy ($\Delta$G) analyses. (**A**) Data obtained for PSS (solid lines) compared to same sequences after they were shuffled (dashed lines). (**B**) Data for randomly picked genomic positions and the associated minimal folding energy. The same number of sites were picked as detected in each strain after the heat shock. Minimal folding energy was calculated at each nucleotide position using a sliding window of 25 nt and a step size of 1 nt for a region of 150 nt up- and downstream of the indicated positions.

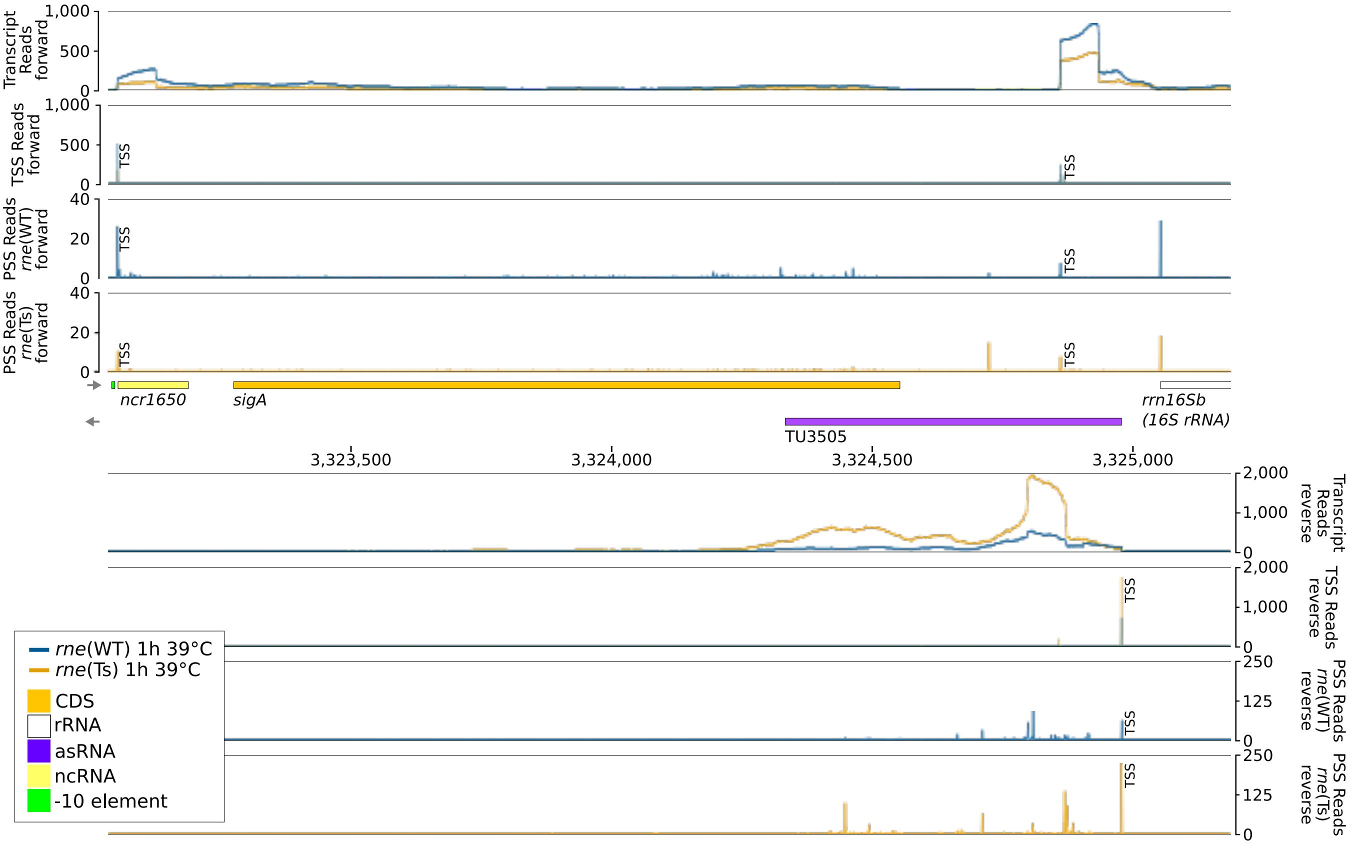


**Supplementary Figure 11.** Genomic locus of operon of *ncr1650* and *sigA*, asRNA to *sigA* (TU3505) and beginning of 16S rRNA. Transcriptome coverage, PSS and TSS for forward and reverse strand are given above and below the middle part of the figure, which indicates the localisation of different features, respectively. The transcriptome coverage for the two indicated strains after incubation for 1 h at 39°C is given on top of the four lanes. Cleavage sites are displayed in the diagrams underneath by the blue and orange peaks, representing PSS detected in *rne*(WT) and *rne*(Ts), respectively. Transcriptome coverage and cleavage sites (PSS) represent the average of normalised read counts of the three investigated replicates. For visualisation, multi-mapping reads were included. Positions which were classified as TSS using DESeq2 are labelled with "TSS" next to the respective peaks. CDS: coding sequences.

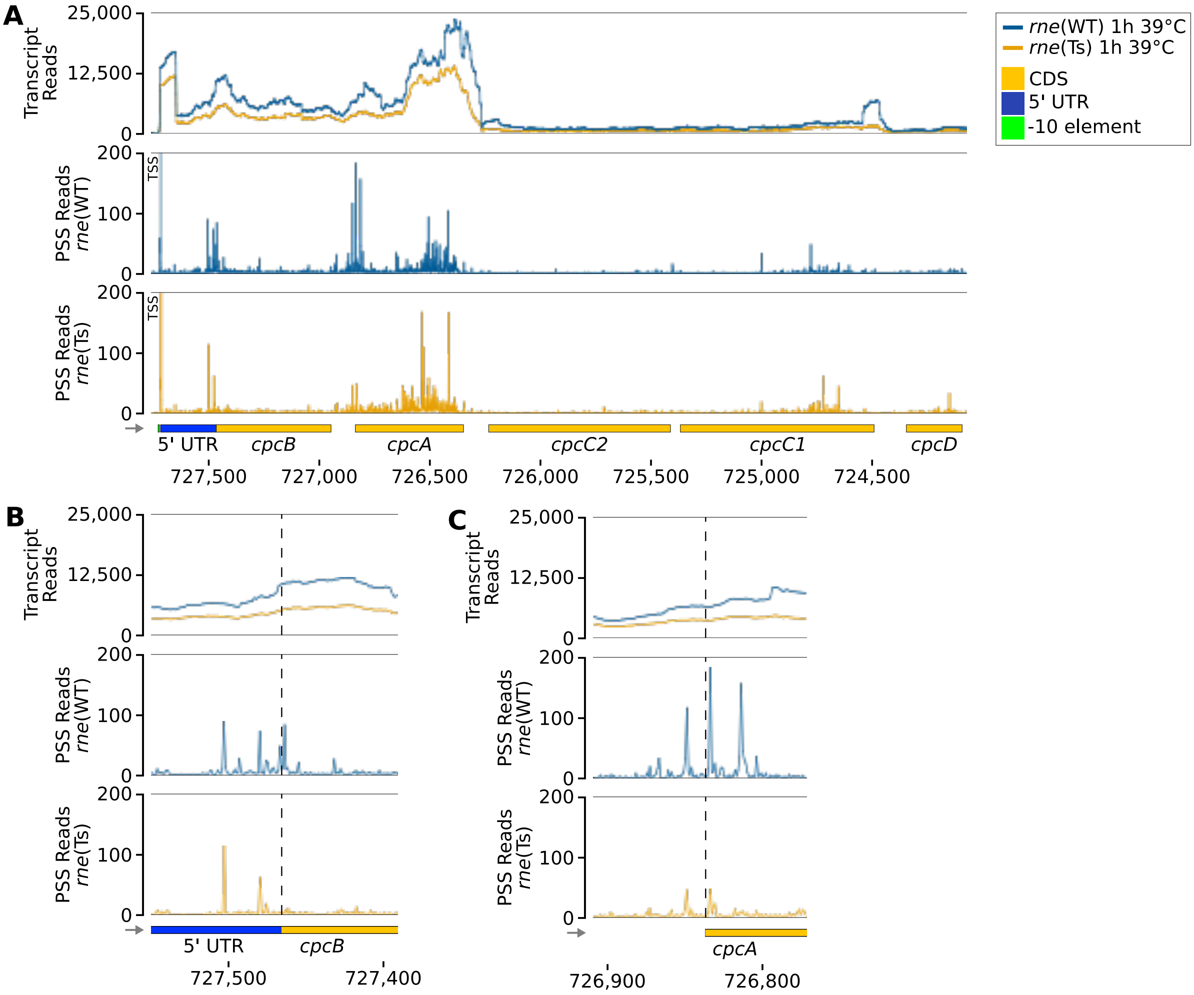


**Supplementary Figure 12.** RNase-E-dependent processing events of the *cpcBAC2C1D* transcript. (**A**) Overview of the *cpcBAC2C1D* operon. (**B**) The region around the start codon of *cpcB*. (**C**) The region around the start codon of *cpcA*. The transcriptome coverage for the two indicated strains after incubation for 1 h at 39°C is given on top of the three lanes. Cleavage sites are displayed in the diagrams underneath by the blue and orange peaks, representing 5'-P (processing sites, PSS) detected in *rne*(WT) and *rne*(Ts), respectively. Transcriptome coverage and cleavage sites (PSS) represent the average of normalised read counts of the three investigated replicates. Positions which were classified as TSS using DESeq2 are labelled with “TSS” next to the respective peaks. Dashed lines in (B) and (C) indicate the start of the coding sequence of *cpcB* and *cpcA*, respectively. CDS: coding sequences.
